# Over-expression and increased copy numbers of a cytochrome P450 and two UDP-glucuronosyltransferase genes in macrocyclic lactone resistant *Psoroptes ovis* of cattle

**DOI:** 10.1101/2025.02.06.636820

**Authors:** Jack Hearn, Wouter van Mol, Roel Meyermans, Kathryn Bartley, Tyler Alioto, Jessica Gomez-Garrido, Fernando Cruz, Francisco Câmara Ferreira, Marta Gut, Ivo Gut, Nadine Buys, Steven Janssens, Karyn Adams, Sara Roose, Thomas Van Leeuwen, Wannes Dermauw, John S. Gilleard, Russell Avramenko, Peter Geldhof, Edwin Claerebout, Stewart T.G. Burgess

## Abstract

*Psoroptes ovis* is a mite species that feeds on sheep, cattle, other ungulates, rabbits, and horses, which can develop into a severe exudative dermatitis known as psoroptic mange. The macrocyclic lactone (ML) family of acaricides are commonly used to control psoroptic mange. However, certain strains of cattle and sheep mites have developed resistance against MLs, which has led to reduced treatment efficacy and even treatment failure.

Here we investigated the genetic basis of ML resistance in mites collected from cattle across Belgium. We compared gene expression between susceptible and resistant mites before and after exposure to ivermectin and genetic diversity between a single susceptible and resistant populations. We generated chromosomal genome assemblies of *Psoroptes ovis* derived from sheep and cattle respectively and correlated genomic diversity of susceptible and resistant mite populations sampled across Belgium.

Gene expression data revealed constitutive over-expression of a cytochrome P450 monooxygenase (CYP) gene and two tandemly located UDP-glucuronosyltransferase (UGT) genes among others. On investigation of the genomic data, we observed copy number variation at both loci in population genomic data. The CYP gene is not amplified in the susceptible population but occurs in multiple copies in all resistant populations and is associated with a peak in F_st_ between resistant and susceptible populations indicative of selection. By contrast, the two UGT genes are massively and tandemly amplified in all populations including the susceptible dataset with a weaker associated signal of selection than the amplified CYP. Hence, distinct mechanisms of amplification and gene regulation are occurring at these putative resistance loci in *P. ovis*.

**Author Summary:** For the first time, we investigated the genetic basis of resistance in scab mites to a key family of drugs (Macrocyclic Lactones) used to control this parasite. Scab mite infestations can cause serious welfare issues in affected cattle and sheep with associated economic impact on production. To identify genes associated with resistance, we applied a combination of approaches including targeted sequencing of candidate genes, genome sequencing and gene expression comparisons of resistant mites with mites that are still susceptible to treatment. We were able to rule-out one family of candidate genes but identified very high expression of genes from two gene families that metabolise, and thereby detoxify, the treatment drug in resistant mites. On examination of the genomic context of these genes we found that the genes had undergone an increase in copy number in the resistant mites compared to the susceptible mites. With our now much increased understanding of resistance in scab mites we can track the spread of resistance using markers in these genes present in resistant mites. We can also now test the suitability of alternative treatments that restore the lethality of Macrocyclic Lactones in scab mites, mitigating the damaging effects of resistance in this species.

## 1. Introduction

*Psoroptes ovis* (Family: Psoroptidae) is an ectoparasitic mite (Class: Arachnida) that feeds on sheep, cattle, other ungulates, rabbits, and horses. Characterised by severe itching, wool loss, and open wounds resulting from self-excoriation, the disease inflicts significant distress on affected animals, leading to substantial economic losses estimated at £80-200 million annually for the UK sheep industry alone [1]. In beef cattle, infestations with *P. ovis* can develop into a severe exudative dermatitis known as psoroptic mange. The severity of the subsequent clinical signs of psoroptic mange differs between individual animals and breeds. Breeds such as the Belgian Blue cattle often develop severe clinical signs that cannot be controlled without the use of acaricides [2–6]. However, certain strains of cattle and sheep mites have developed resistance against the commonly-used macrocyclic lactone (ML) family of acaricides, which can lead to reduced treatment efficacy and even treatment failure [7–13]. Unfortunately, insights into the underlying resistance mechanisms of *P. ovis* against MLs and other acaricides are lacking.

Based on knowledge from the two-spotted spider mite, *Tetranychus urticae*, in which acaracide resistance mechanisms have been well studied [14], acaricide resistance in mites likely develops via two main mechanisms: 1) a pharmacokinetic mechanism, mainly implemented through changes in detoxification enzymes and channel transporters; 2) a pharmacodynamic mechanism, involving decreases in drug sensitivity due to target site changes [15,16]. Such metabolic and target-site mechanisms of resistance have rapidly evolved in many arthropod vector and pest species under selection pressure from pesticide use including various fly species, aphids, and beetles among others [14,17]. Underlying genetic changes often differ between these two mechanisms with single non-synonymous mutations capable of conferring target-site resistance. Metabolic resistance may result from mutations that affect the promoter regions (cis acting) and regulators (trans) of genes with subsequent change in gene expression levels and gene duplication or deletion events that result in copy number changes of key loci [18,19]. Combinations of these mechanisms have also been found [14,20]. Furthermore, mechanisms that result in gene overexpression may act in tandem with coding-sequence mutations that enhance the affinity of a protein for a specific pesticide. The target site of MLs is the family of cys-loop ligand-gated ion channels (cysLGIC) found in vertebrates and invertebrates [21]. A functional cysLGIC consists of 5 subunits and each subunit typically has 4 transmembrane regions and a conserved disulphide bridge motive at the extracellular domain of the N-terminus. Glutamate, γ-aminobutyric acid (GABA), histamine and acetylcholine act as ligands, as well as shifts in extracellular proton concentration [21,22]. cysLGICs are distributed throughout the nervous system of invertebrates and have inhibitory (anion) and excitatory (cation) functions [23]. CysLGIC candidate genes activated by ML binding, specifically ivermectin, include Glutamate-gated chloride channels (GluCl), GABA-receptors (GABA-Cl), histamine-gated chloride channels (HisCl) and pH-sensitive chloride channels (pH-Cl) [24–26]. Mutations in GluCls, mostly in the third trans membranal (TM) region, have been linked to ML resistance in a number of arthropods [21,27]. GluCls are expressed in sensory neurons, interneurons and motoneurons and play a role in a considerable number of functional behaviours, e.g. pharyngeal pumping, frequency of change in movement direction and heat and odour responses [22,28–30]. From all cysLGICs, the MLs have the highest affinity for the GluCls [31]. Histamine is the predominant neurotransmitter of arthropod photoreceptors, and consequently HisCls have been observed in arthropod eyes. They also play a role in temperature tolerance and preferences in *D. melanogaster* [32]. They are susceptible to avermectins and may play a role in the neurotoxic effects of the MLs [33,34].

Pharmacokinetic or metabolic changes can decrease the bioavailability of a xenobiotic in three possible phases as part of the detoxification pathway. In Phase I, cytochrome P450 monooxygenases or esterases increase the polarity and reactivity of the xenobiotic through the addition of a hydroxyl, carboxyl or amino group. Phase II enzymes conjugate glutathione by glutathione S-transferase or urine diphosphate (UDP) by UDP-glycosyltransferase to either the xenobiotic or a Phase I product. A UGT gene confers resistance to the ML abamectin in the citrus mite *Panonychus citri* [35]. In Phase III, these polar metabolites or the xenobiotic itself are transported away from target cells by the ATP-binding cassette transporters (ABC-transporters) [15,16]. Many of the genes in this pathway across all three phases of the detoxification pathway have been implicated in resistance in many arthropods, with examples from other species of mite [14]

Other known mechanisms of resistance include changes to behaviours or arthropod cuticles. Cuticular resistance acts by preventing, or slowing, the penetration of a pesticide to its target site through modification of the chitin proteins and complex hydrocarbon mix that make-up the cuticle. Behavioural resistance results from a modification in behaviour that reduces pesticide exposure and has been observed in *T. urticae* mites [36,37]. The genetic basis of behavioural resistance is the least well understood of all the known mechanisms of pesticide resistance among arthropods and it is possibly of more importance in flying arthropods than mites.

The first objective of this study was to identify the cysLGICs in *P. ovis* and to explore possible target site variations in candidate GluCl genes from multiple *P. ovis* isolates with different ML susceptibility. The second objective of this study was to identify potential resistance mechanisms for macrocyclic lactones in *P. ovis,* by contrasting and intersecting gene expression and genomic signals of selection from ML susceptible and ML resistant cattle mite populations sampled in Belgium. In this case, the transcriptome response of mites from the resistant population was studied before and after exposure to the ML to define differences in constitutive versus induced gene expression. Both ML-exposed and unexposed resistant mites were contrasted against the unexposed susceptible population and versus one another. Genomic data was collected from multiple populations of cattle mites across Belgium and susceptibility to MLs was assayed per site. Multiple individuals were subsequently combined post-exposure for pooled-template whole genome sequencing (PoolSeq). These pooled data were contrasted between susceptible and resistant populations to identify regions of strong differentiation. Candidate loci were subsequently investigated for underlying genetic changes which correlated with observed phenotypes.

## 2. Material and methods

### 2.1. Target site variation in cys-loop ligand-gated ion-channels

#### 2.1.1. cys-loop ligand-gated ion-channel identification in *P. ovis*

The Online Resource for Community Annotation of Eukaryotes (OrcAE; https://bioinformatics.psb.ugent.be/orcae/overview/Psovi) database was tBLASTn-searched for genes encoding for cysLGIC in the current *P. ovis* reference genome [38]. Known protein sequences of GluCls, GABA-Cl, pH-Cl, HisCl and nAchR from the two-spotted spider mite, *Tetranychus urticae*, the fruit fly, *Drosophila melanogaster*, the scabies mite, *Sarcoptes scabiei*, the tick, *Rhipicephalus microplus* and the house dust mite, *Dermatophagoides pteronyssinus* were used to screen the genome for homologues (e-value threshold of < 1e-50). These protein sequences were extracted from the database of the National Center for Biotechnology Information (https://www.ncbi.nlm.nih.gov/). The most likely cysLGIC subunit encoding genes from *P. ovis* were identified based on their homology at the amino acid level.

Information on the transcription levels of the cysLGIC subunit encoding genes during the life cycle of *P. ovis* was extracted from OrcAE using lifecycle stage-specific gene expression data [39]. A heatmap was constructed with the transcription data of the genes across the different lifecycle stages, i.e. larvae, protonymphs, tritonymphs, adult females and adult males, from [39] in R (R Core Team, 2020). CysLGIC subunits from *P. ovis, T. urticae, D. melanogaster, S. scabiei* and *D. pteronyssinus* were aligned with the use of MUSCLE [41]. The Jones, Taylor and Thornton model was used for the phylogenetic analysis. A maximum likelihood analysis, bootstrapping 1000 pseudo-replicates, was performed with MEGA X [42] to construct a midpoint rooted tree.

#### 2.1.2. Sample collection

Pre-treatment skin scrapings from 9 farms corresponding to an ML field efficacy study by [13] were used for the collection of 50-100 living *P. ovis* mites per farm (Table 1). Mite isolates from the different farms differed in their susceptibility to ML treatment (ivermectin, doramectin and moxidectin), as determined by the calculation of the mite count reduction two weeks post-treatment. Mite containing skin samples from sheep were collected from infested donor animals held at the Moredun Research Institute, UK. These mites have never been exposed to acaricides throughout their maintenance as a reference population and are therefore completely susceptible to treatment with moxidectin (S. Burgess, Personal Communication). Before collection, skin scrapings and samples were incubated for 20 minutes at 37°C to increase mobility of the mites. Samples were screened with a stereomicroscope and viable mites from all lifecycle stages were collected with a needle. 25 mites per sample were snap frozen in liquid nitrogen and stored at -80°C.

**Table 1.** Overview of the different samples used for DNA extraction together with the number of mites, region of origin, in vivo macrocyclic lactone susceptibility and host species as established by [13].

| Sample name | No. mites | Region of origin | Macrocyclic lactone susceptibility | Host |
| --- | --- | --- | --- | --- |
| D03 | 75 | West-Flanders, Belgium | Susceptible | Belgian Blue Cattle |
| D05 | 75 | East-Flanders, Belgium | Suspected Resistant | Belgian Blue Cattle |
| D06 | 75 | East-Flanders, Belgium | Resistant | Belgian Blue Cattle |
| D07 | 75 | East-Flanders, Belgium | Suspected Resistant | Belgian Blue Cattle |
| D08 | 75 | East-Flanders, Belgium | Suspected Resistant | Belgian Blue Cattle |
| D09 | 50 | Zeeland, The Netherlands | Resistant | Belgian Blue Cattle |
| D10 | 50 | West-Flanders,<br>Belgium | Resistant | Belgian Blue Cattle |

#### 2.1.3. DNA extraction

The temperature of a pre-chilled mortar (overnight at -80°C) was further lowered with liquid nitrogen. When most nitrogen had evaporated, 25 previously snap-frozen mites were submerged in a thin layer of liquid nitrogen. ATL-buffer (180 µl per 25 mites, Qiagen^®^, DE) was added when all the liquid nitrogen had evaporated. The mites were crushed with a pestle to a fine powder and transferred to a new 1.5 ml eppendorf tube. To each sample, 20 µl proteinase K (Qiagen®, DE) was added and the solution was incubated overnight in a shaking heat block at 56°C at 300 RPM. Five hundred µl phenol:chloroform:IAA (25:24:1; Sigma-Aldrich, USA) was added. The mixture was briefly vortexed and centrifuged for 10 minutes at 10,000 g (4°C). The supernatant (≈ 400 µl) was transferred to a new 1.5 ml eppendorf tube and 40 µl (=1/10) sodium acetate (3M; pH 5.2; Sigma-Aldrich, USA) and 400 µl (=1/1) isopropanol (Biosolve, NL) were added. After centrifuging for 10 minutes at 16,000 g (4°C), the supernatant was discarded.

Six hundred µl 80% ethanol (Sigma-Aldrich, USA) was added to the pellet and centrifuged for 5 minutes at 10,000 g (4°C). After discarding the supernatant, this washing step was repeated once. Finally, the pellet was dried at room temperature (∼20°C) until the ethanol had evaporated (∼5 to 15 minutes) and subsequently dissolved in 25 µl nuclease-free water (Ambion, USA). The DNA concentration was measured with a NanoDrop™ 2000 spectrophotometer (Thermo Fisher Scientific, USA).

#### 2.1.4. High fidelity PCR and purification

Fragments (179 bp long) from the two different GluCls in *P. ovis* were amplified according to the Phusion^®^ High-Fidelity polymerase chain reaction protocol (Thermo Fisher, USA). The forward primer for GluCl-44 was 5’ AGCCGCCTGTCAGTTATACG 3’ and the reverse primer 5’ GCAATCCATAAGCAGCCAAT 3’. For GluCl-280, the forward primer was 5’ ACATCCATCAGTGGCATCAA 3’ and the reverse primer 5’ GGCTGAACCATGTTCGAGAT 3’ (detailed in Supplementary Table S1. The thermocycling parameters were 98°C for 30 s, followed by 30 cycles of 98°C for 5 s, 62°C for 10 s, 72°C for 15 s, and a final extension of 72°C for 5 min. PCR products were purified from agarose gels with the Zymoclean Gel DNA Recovery kit (Zymo Research, USA), following the manufacturers’ protocol.

#### 2.1.5. Deep amplicon sequencing

##### 2.1.5.1. PCR amplification of the GluCl-44 and GluCl-280 fragments

Locus specific primers were designed to amplify a 219 bp fragment of interest from both GluCl-44 (PsoOvis1B009037 in CNAG genome annotation described below) and GluCl-280 (PsoOvis1B011928). These primers were adjusted for Illumina deep sequencing and for both genes, four forward and four reverse primers were mixed in equal concentrations (primer design, Supplementary Table S1). The Illumina adaptor oligonucleotide sequences were obtained from the Illumina Adapter Sequences document (Oligonucleotide sequences^©^ 2020 Illumina, Inc.) and added to allow later annealing of sequencing primers. For each of the primers, 0-3 random nucleotides were inserted between the locus specific primer sequence and the Illumina Adapter sequence. These nucleotides prevent oversaturation of the MiSeq sequencing channels by increasing the diversity of generated amplicons. PCR conditions as follows: 5 μL KAPA HiFi HotStart Fidelity Buffer (5X) (KAPA Biosystems, USA), 0.75 μL Forward Primer Mix (10 μM), 0.75 μL Reverse Primer Mix (10 μM), 0.5 μL dNTPs (10 mM), 0.5 μL KAPA HiFi HotStart Polymerase (0.5 U), 16.4 μL molecular-grade H_2_O, 0.1 μL BSA (20mg/mL) and 1μL of 1:10 dilution of gel-purified PCR product. The thermocycling parameters were 95°C for 3 min, followed by 30 cycles of 98°C for 10s, 65°C for 15s, 72°C for 15s, and a final extension of 72°C for 2 min. PCR products were purified with AMPure XP Magnetic Beads (1X) (Beckman Coulter, Inc., USA) following the manufacturer’s recommended protocol. All samples were eluted in 32.5 μL of molecular-grade H_2_O.

##### 2.1.5.2. Addition of indices, pooling and sequencing of the library

By limited cycle PCR amplification, both Illumina barcode indices and P5/P7 sequencing regions were added to the GluCl-44 and GluCl-280 amplicons. Unique forward/reverse combinations of primers of the Nextera XT Index Kit v2 set (Oligonucleotide sequences^©^ 2020 Illumina, Inc.) were made, enabling individual sample barcoding. The following PCR conditions were used: 5 μL KAPA HiFi HotStart Fidelity Buffer (5X) (KAPA Biosystems, USA), 2.5 μL Forward Primer (S502-S511) (5 μM), 2.5 μL Reverse Primer (N716-N719) (5 μM), 0.75 μL dNTPs (10 mM), 0.5 μL KAPA HiFi HotStart Polymerase (0.5 U), 8.75 μL molecular-grade H_2_O and 5μL of first-round clean PCR product as template. The thermocycling parameters were 98°C for 45s, followed by seven cycles of 98°C for 20s, 63°C for 20s, 72°C for 2 min. Amplicons were purified with AMPure XP Magnetic Beads (1X) (Beckman Coulter, Inc., USA) as previously described.

The concentration of the second-round clean PCR product was measured using the Implen (DE) NanoPhotometer^®^ NP80 and 50ng of each sample was pooled to produce a master sequencing library. The final concentration of this pooled library was assessed with the KAPA qPCR Library Quantification Kit (KAPA Biosystems, USA), following the manufacturer’s recommended protocol. The prepared pooled library was run on an Illumina MiSeq Desktop Sequencer using a 2x250 v2 reagent kit (MiSeq Reagent Kits v2, MS-103-2003) at a concentration of 15 pM with the addition of 20% PhiX Control v3 (Illumina, FC-110-3001). The MiSeq was set to generate only FASTQ files with no post-run analysis. Based on their supplied index combinations, samples were automatically demultiplexed by the MiSeq. All protocols were carried out per Illumina’s standard MiSeq operating protocol (Illumina, Inc., USA).

Raw FASTQ files generated were analyzed with the DADA2 v.1.11.5 bioinformatic software package to ascertain the number of unique amplicon sequencing variants (ASV) contained in each sample (PMID: 27214047). This workflow was adapted from the DADA2 analysis described at www.nemabiome.ca, with modifications to accommodate for the amplicons analyzed in this paper. Briefly, FASTQ files were pre-filtered with the ‘FilterAndTrim’ function to remove any ‘Ns contained in the sequences. Cutadapt was used to remove forward and reverse primer sequences in the amplicons (PMID: 28715235). After primer removal, reads were filtered again with ‘FilterAndTrim’ with no N’s allowed, maxEE = 6, truncQ = 2, a minimum length of 200 bp for each forward and reverse read, and the removal of phiX if identified. A sequence table was constructed with ‘makeSequenceTable’ to display all ASVs present in the dataset. Chimeras were removed with ‘removeBimeraDenovo’. This provides a read count of each ASV present in each sample. A fasta file was also generated with the ‘getUniques’ and ‘uniquesToFasta’ commands to provide a list of all ASVs identified and their corresponding nucleotide sequence. Each ASV was blasted against a reference sequence to ensure that the ASV correctly matched the intended amplicon. Any off-target amplicons were subsequently removed from analysis. Furthermore, ASVs with extremely low reads were manually deleted from the dataset since these most likely represent PCR artifacts. All amplicon sequences were submitted to the European Nucleotide Archive (ENA) under PRJEB82994.

### 2.2 Chromosomal sheep-derived *Psoroptes ovis* genome and guided-assembly of cattle *P. ovis* genome

To address fragmentation with the previous iteration of the *P. ovis* genome, which contained 93 contigs [38], a new *P. ovis* sheep mite-derived genome assembled into chromosomes was generated A *P. ovis* cattle mite genome was subsequently assembled using the *P. ovis* sheep scab genome as a reference. Sequencing and genome assembly and annotation were undertaken by the Centro Nacional de Análisis Genómico (CNAG, Spain) using their assembly pipeline (https://github.com/cnag-aat/assembly_pipeline).

Identification of cys-loop ligand-gated ion-channels and qPCR analyses were undertaken using an earlier version of the *P. ovis* geneset as a basis [39]. A BLAST-based reciprocal best hits (RBH) approach [43] was used to identify corresponding genes between our newly generated geneset and the geneset of [39]. BLAST searches were performed at the protein level. We use gene names from our CNAG-generated dataset throughout the text to avoid confusion and present corresponding gene names for both genesets in tables.

#### 2.2.1. Long-Read Sequencing

Genomic DNA from *P. ovis* isolated from sheep (UK) and from cattle (Belgium) was extracted from adult female *P. ovis* mites and quality-controlled for long-read sequencing, considering purity, quantity and integrity. The sequencing libraries were prepared using the ligation 1D sequencing kit 9 or upgraded kit 10 chemistry (SQK-LSK109 or SQK-LSK110, respectively) from Oxford Nanopore Technologies (ONT). In brief, 3.0 μg of the gDNA underwent end-repair and adenylation using the NEBNext UltraII End Repair/dA-Tailing Module (NEB), followed by ligation of sequencing adaptors. The ligation product was purified using 0.4X AMPure XP Beads and eluted in Elution Buffer (ONT). The WGS long-read sequencing run was performed on a GridIon Mk1 (ONT) using a flow cell R9.4.1 (FLO-MIN106) compatible with sequencing ligation kit 9 and 10. The sequencing data was collected for 72 hours. The quality parameters of the sequencing runs were monitored in real time using the MinKNOW platform version 4.2.5, and the basecalling was performed using Guppy version 4.3.4.

#### 2.2.2. Short-Read Whole Genome Sequencing (WGS) Library Preparation

The short-insert paired-end library for whole genome sequencing was prepared using the PCR-free protocol and the KAPA HyperPrep kit (Roche). After end-repair and adenylation, Illumina platform-compatible adaptors with unique dual indexes and unique molecular identifiers (Integrated DNA Technologies) were ligated. The sequencing libraries were quality controlled on an Agilent 2100 Bioanalyzer using the DNA 7500 assay (Agilent) to assess size and quantified using the Kapa Library Quantification Kit for Illumina platforms (Roche).

#### 2.2.3. 10X Genomics library preparation and short read sequencing

The linked read libraries from each isolate of gDNA were prepared using the Chromium Controller instrument (10x Genomics) and Genome Reagent Kits v2 (10x Genomics) following the manufacturer’s protocol. Briefly, 10 ng of high molecular weight genomic DNA (HMW gDNA) was portioned in GEM reactions, including a unique barcode (Gemcode), after being loaded onto a chromium controller chip. The droplets were then recovered, isothermally incubated, and fractured, and the intermediate DNA library was then purified and size-selected using Silane and Solid Phase Reverse Immobilisation (SPRI) beads. Illumina-compatible paired-end sequencing libraries were prepared following 10X Genomics recommendations and validated on an Agilent 2100 BioAnalyzer with the DNA 7500 assay (Agilent). The 10X Genomics and WGS libraries were sequenced on Illumina NovaSeq 6000 with a read length of 2x151bp, following the manufacturer’s protocol for dual indexing. Image analysis, base calling, and quality scoring of the run were processed using the manufacturer’s software, Real Time Analysis (RTA 3.4.4). Read sequences and assemblies for the sheep-derived and cattle-derived *P. ovis* assemblies are available under ENA project PRJEB84953 in sub-projects PRJEB82899 and PRJEB82900 respectively).

#### 2.2.4. Genome assembly and annotation

For the sheep-derived assembly, filtered long read (ONT) data was assembled with Flye v2.8.3 [44] with two iterations of polishing. To improve the base accuracy of the assembly, it was polished three times with HyPo [45] using both Illumina and ONT data. To remove potential contamination, we examined read coverage (ONT reads were aligned back to the assembly with minimap2 [46]) as well as taxonomic identification by searching against the NCBI nt database using megablast. Only those contigs with hits to mite sequences and with the expected read coverage were retained. The decontaminated assembly was scaffolded with 10X Linked-Reads using the Faircloth’s Lab pipeline (http://protocols.faircloth-lab.org/en/latest/protocols-computer/assembly/assembly-scaffolding-with-arks-and-links.html#). For the cattle-derived *P. ovis* data, the genome was assembled using the CLAWS v2.1 workflow [47] combining ONT long reads and 10X linked reads. As this assembly was of low contiguity, the assembly was then aligned to the sheep-derived assembly and reference-based scaffolding was carried out using RagTag [48]. We provide more in-depth methods of assembly, removal of contaminant sequences and assembly evaluation in the Supplementary Methods.

The gene annotation for the sheep-derived *Psoroptes ovis* genome assembly was obtained by combining transcript alignments of available *P. ovis* RNASeq data, protein alignments of related species and *ab initio* gene predictions principally using PASA [49]. Finally, the *P. ovis* sheep genome annotations were mapped to the *P. ovis* cattle genome using Liftoff [50]. A more detailed explanation of genome annotation is given in the Supplementary Methods.

#### 2.2.5. Genome re-sequencing and population genomics

Raw data for the whole genome re-sequencing was deposited in the ENA under accession PRJEB84953. Reads for the seven samples of susceptible, resistant and suspected resistant populations were quality and adapter trimmed with fastp (version 0.23.4) default parameters and sequence quality checked with FastQC (v0.12.1) [51]. Trimmed reads for each population were aligned to the sheep and cattle *P. ovis* genomes separately with BWA (v0.7.17) [52]. PCR duplicates of aligned reads were marked and subsequently sorted by mapping coordinates using Picard (v2.18.29) [53]. Alignment statistics for each processed BAM file were generated using samtools “flagstats” and “stats” tools (v1.17) [54]. Variant predictions across populations were made in Varscan (v2.4.6) [55] as it is compatible with pooled template sequencing by estimating allele frequency per population. Samtools mpileup [54] was used to generate the Varscan input format and a p-value threshold of <0.05 set and a minimum variant frequency of 0.01 required for variant prediction. The gene body location and potential effect of variants was predicted in SnpEff (v5.2) [56] using a custom database for the sheep and cattle genomes. Average coverage per gene was calculated with the bamstats05 tool of the Jvarkit package (dbdbed3a9) [57]. Average read coverage for each gene across the sheep and cattle genomes was divided by median coverage for all genes in order to identify outlier loci and coverage plots created in the R package ggplot2 (v3.4.4). Fixation indices estimated by the unbiased-hudson method (F_st_) were calculated from bam files per gene and in sliding windows of 10,000 bp with 1,000 bp steps using the Grenedalf package (v0.3.0) [58]. Grenedalf is a re-implementation of the popoolation2 package [59]. Tajima’s and nucleotide diversity (π) were also estimated but discarded due to low pool coverages resulting in many immeasurable regions of the genome. Poolfstat (v2.2.0) [60] and pcadapt [61] were used to generate global pairwise nucleotide distance and F_st_ estimates and perform principal components analysis (PCA) between populations respectively. Smoove (v.0.2.8) (https://github.com/brentp/smoove) was used to predict and annotate structural variants.

Regions of interest in the genome predicted computationally were visually checked using the Integrative Genomics Viewer (v2.16.2) (IGV) [62]. Oxford nanopore reads sequenced for cattle and sheep genome assemblies were revisited to identify the genomic context of genes of increased copy number among resistant and susceptible populations. Protein sequences of genes with elevated coverage were searched against Oxford Nanopore reads with Diamond (v2.1.8) [63] to determine if tandem duplications (cis) or insertions in other regions of the genome (trans) could explain observed increases in copy number. Diamond was run in ultra-sensitive mode with the number of query nanopore reads to subject *P. ovis* protein sequences increased to 1000 to accommodate more than one copy of a gene occurring discretely on a single read. In order to enrich the results for the most informative reads, a minimum of 10 hits (HSPs) per read were considered for investigating inflated tandem copy numbers.

### 2.3. Gene expression differences in ML resistant and susceptible *P. ovis*

Through RNA sequencing and validation with Real Time quantitative PCR (qPCR), the transcriptomes of a ML susceptible mite population (SUS), a ML-resistant mite population before exposure to ivermectin (RES_unexposed_) and the same ML-resistant mite population after exposure to ivermectin (RES_exposed_) were compared. Each analysis was performed in triplicate as this degree of replication had previously been demonstrated to be sufficient to identify significant changes in gene expression at a level of fold-change >2 in *P. ovis* [39].

#### 2.3.1. Sample collection

A farm with a known ML susceptible *P. ovis* population, 99.3% mean reduction in mite counts post-treatment, and six assessed as resistant mite population with a mean reduction of 36% in mite counts post-treatment from the field efficacy study [13] were revisited to collect susceptible and resistant mites before and after treatment, (see Table 1). On both farms, a group of 12 animals with clinical psoroptic mange was selected and physically separated from the rest of the herd. From this group, 6 animals were selected for validation of the ML susceptibility at the time of sampling as described in [13], and 6 for the collection of mites for RNA extraction. The pre-treatment samples were taken from a maximum of half of the lesion surfaces. All animals were subsequently treated twice with a subcutaneous injection of Ivomec (Merial, FR) at a dose of 0.2 mg ivermectin per kg with a 7-day interval. On the ‘resistant’ farm, the post-treatment samples were collected 7 days after the second ivermectin treatment. All samples were processed within 8 hours after collection. After incubation at 37°C, adult female *P. ovis* mites were collected from the skin scrapings with a fine needle. Differentiation of life-cycle stages was based on [64]. Mites were snap-frozen in liquid nitrogen in batches of 25 and stored at -80°C in 2 ml tubes (VWR™, USA).

#### 2.3.2. RNA extraction and quality control

A glass homogenizer (7 ml, Kontes^®^, USA) was soaked in 0.1% DEPC for 12 hours at room temperature (∼20 °C). A total of 150 frozen adult female mites were put into the glass homogenizer and 1 ml TRIZol (Ambion, USA) was added before homogenisation. Samples were homogenised until all macroscopically visible particles in the solution were gone. One ml of the homogenised mixture was transferred into a 2 ml tube and incubated for 5 minutes at room temperature, (∼20 °C). The samples were then centrifuged at 12,000 g for 10 minutes in a pre-chilled rotor (4°C) and the supernatant transferred to a new 2 ml tube. Two hundred µl chloroform (Sigma-Aldrich, USA) was added to each tube and the tubes were inverted end-on-end 10 times. The samples were then centrifuged at 10,000 g (4°C) for 15 minutes and the top aqueous phase carefully transferred to fresh 2 ml tubes. Five hundred µl isopropanol (2-propanol; Biosolve, NL) was added to each aqueous phase sample and the tubes were inverted end on end 10 times. The samples were then centrifuged at 12,000 g (4°C) for 15 minutes and the supernatant discarded. One ml of 75% ethanol (Sigma-Aldrich, USA) was added to each pellet and then mixed by vortexing. The samples were subsequently centrifuged at 7,500 g (4°C) for 5 minutes. The ethanol was decanted and the pellets were air dried for 1-2 minutes at room temperature (∼20 °C). The RNA-pellet was resuspended in 50 µl RNase free water (Ambion, USA) and stored at -80°C. RNA quantity was determined with the NanoDrop 2000 (Thermo Fisher Scientific, USA). RNA quality was determined with the Experion™ Automated Electrophoresis Station (Bio-Rad, USA).

#### 2.3.3. RNA seq

All 9 replicates were diluted to a final concentration of 40 ng/µl RNA with nuclease free water (Ambion, USA). From the RNA samples, Illumina libraries were constructed with the TruSeq Stranded mRNA Library Preparation Kit (Illumina, USA), and the resulting libraries were sequenced on a NovaSeq 6000 to generate strand-specific paired reads of 2 × 100 bp. Library construction and sequencing was performed at Fasteris, Geneva, Switzerland and raw reads are available under project PRJEB82994 in the ENA. Post-sequencing, read quality of raw FASTQ files was checked with FastQC v0.11.9. Pseudo alignment of the RNA-seq reads to the CNAG *P. ovis* sheep scab transcriptome was performed using Kallisto (v1.15.0) [65] as part of the rna-seq-pop package (v1.0.4) [66]. Kallisto generated read counts for all RNA-seq samples were used as input for DESeq2 (v1.30.1) [67] and Sleuth [68] to identify significantly differentially expressed genes and transcripts respectively between the 3 ML-resistant unexposed (RESunexposed), 3 ML-resistant exposed (RESexposed) and 3 ML-susceptible (SUS) replicates. Significantly differentially expressed genes and transcripts were classified as those having a fold change ≥ 2.0 between each of the pairwise comparisons and a False Discovery Rate (FDR) corrected p-value of ≤ 0.05.

#### 2.3.4. Validation of RNA-seq data by Real Time qPCR

qPCR was used to validate the transcription of genes associated with ML resistance in the tested *P. ovis* populations. qPCR was performed on cDNA samples from all nine RNA samples used for the original RNA-seq analysis SUS x3, RES_unexposed_ x3, RES_exposed_ x3 and six differentially expressed genes were selected along with three *P. ovis* housekeeping genes. cDNA libraries were constructed with the iScript cDNA Synthesis Kit (Bio-Rad Laboratories Inc., USA) following the recommended protocol. Forward and reverse primers were designed for six genes of interest from the RNASeq experiment using the gene sequences of the Burgess et al., (2018) *P. ovis* genome [38]: two UDP-glycosyltransferases, PsoOvis1B004414 and PsoOvis1B010992, an inositol oxygenase, PsoOvis1B000159, CYP genes PsoOvis1B011549 and PsoOvis1B005763, and a glutathione S-transferase, PsoOvis1B008730, and for three housekeeping genes: a beta actin, PsoOvis1B008887/PsoOvis1B000861 [69], a mitochondrial ribosomal protein, PsoOvis1B008370 [21], and elongation factor 1-alpha, PsoOvis1B001219, [70]. A description of primers used for each gene is given in Supplementary Table S2. All primer sequences were BLAST searched against the *P. ovis* genome to rule out cross specificity.

qPCR reactions were run on a StepOnePlus real-time PCR system (Applied Biosystems, USA). cDNA was diluted 1:8 in nuclease free water and Fast SYBR® Green Master Mix Real-Time PCR Master Mix was used following the recommended protocol. Standard curves for all genes were based on a pooled sample from SUS1, SUS2 and SUS 3 in 1:2, 1:4, 1:8, 1:16 and 1:32 dilution. All samples were run in duplicate. PCR conditions were as follows: 1 cycle of 95°C for 20 s, 40 cycles of 3 s at 95°C and 30 s at 60°C (optimal annealing temperature for all examined genes) and 1 cycle of 15 s at 95°C, 60 s at 60°C and 15 s at 95°C. Relative quantities of gene expression were calculated using the delta Ct method to determine the fold differences in gene transcription levels of the genes in the different mite populations. Housekeeping genes were used for normalisation with Genorm [71]. Correlation between RNAseq and qPCR values was performed using the R package “rmcorr” Shiny application (https://lmarusich.shinyapps.io/shiny_rmcorr/) [72] following [73].

Rmcorr is able to take into account non-independent measures, in our case each gene was assessed three times for each contrast between RES_unexposed_, RES_exposed_ and SUS replicates. This allows us to include all contrasts in a single correlation analysis. Rmcorr uses analysis of covariance (ANCOVA) to statistically adjust for interindividual variability and fits a common slope and variable intercept to all data points, i.e. genes, included.

## 3. Results

### 3.1. Objective 1. Pharmacodynamic changes

#### 3.1.1. Identification of cys-loop ligand-gated ion-channels

In *P. ovis*, 16 different cysLGICs were found, that are activated by either glutamate (n=2), histamine (n=2), GABA (n=4), acetylcholine (n=7) or environmental pH (n=1).

All cysLGICs showed homologies of 63% or more at amino acid level with at least one other cysLGIC from *T. urticae, D. melanogaster, S. scabiei*, *D. pteronyssinus* and *R. microplus.* Moreover, high levels of homology, between 82% and 99%, were observed with certain cysLGICs in other species, most often with *D. pteronyssinus,* probably due to the relatively close homology between these species [74]. An overview of the different cysLGICs in *P. ovis* is given in Table 2. together with the corresponding cysLGIC with the highest homology in *T. urticae, D. melanogaster, S. scabiei, D. pteronyssinus* and *R. microplus*.

**Table 2.** Summary of the cysLGIC encoding genes in *P. ovis* with the corresponding cysLGICs with the highest homology in *T. urticae*, *D. melanogaster*, *S. scabiei*, *D. pteronyssinus* and *R. microplus*. “NI” indicates that a homologue was (N)ot (I)dentified in that species. Italics represent gene pairs with ambiguous reciprocal best hits due to different copy numbers of the locus between Burgess et al., (2018) [38] and our genome annotation: ^1^Reciprocal best hit is PSOVI295g01540; PSOVI295g01520 and PSOVI295g01540 are represented by a single gene, PsoOvis1B003100, in our *P. ovis* genome annotation; 2 Reciprocal best hit is PSOVI01g00020; PSOVI01g00020 and PSOVI286g03370 are represented by a single gene, PsoOvis1B013090, in our *P. ovis* genome annotation dataset.

|  |  |  |  |  |  |  |
| --- | --- | --- | --- | --- | --- | --- |
| *P. ovis* CNAG<br>Gene ID | *Burgess et al.*<br>Gene ID | *T. urticae* | *D. melanogaster* | *S. scabiei* | *D. pteronyssinus* | *R. microplus* |
| Glutamate-gated chloride channels |  |  |  |  |  |  |
| PsoOvis1B009037 | PSOVI44g00510 | 74% GluCl-3 | 63% GluCl | 94% GluCl KPM04243.1 | 99% XP_027194799.1 | NI |
| PsoOvis1B011928 | PSOVI280g05660 | 82% GluCl-5 | 71% GluCl | 67% GluCl KPM04243.1 | 95% XP_027196448.1 | NI |
| **Histamine-gated chloride channels** |  |  |  |  |  |  |
| *PsoOvis1B003100<sup>1</sup>* | *PSOVI295g01520* | 67% HisCl-1 | 65% HisCl-1 | NI | NI | NI |
| PsoOvis1B003100 | PSOVI295g01540 | 67% HisCl-1 | 61% HisCl-1 | NI | NI | NI |
| **pH-sensitive chloride channel** |  |  |  |  |  |  |
| PsoOvis1B000735 | PSOVI280g03190 | 63% pH-Cl | 55% pH-Cl [25] | 29% pH-Cl | NI | NI |
| **GABA-gated chloride channels** |  |  |  |  |  |  |
| *PsoOvis1B013090<sup>2</sup>* | *PSOVI286g03370* | 71% RDL-2 | 72% RDL | NI | 89% XM_027338589.1 | 58% ACV07674.1 |
| PsoOvis1B013090 | PSOVI01g00020 | 71% RDL-2 | 72% RDL | NI | 89% XM_027338589.1 | 58% ACV07674.1 |
| PsoOvis1B001590 | PSOVI14g08260 | 46% RDL-1 | 84% Lcch3 | NI | 72% XM_027347239.1 | 48% ACV07674.1 |
| PsoOvis1B009531 | PSOVI36g00450 | 77% RDL-2 | 78% RDL | NI | 94% XM_027350548.1 | 79% ACV07674.1 |
| **Nicotinic acetylcholine-receptor** |  |  |  |  |  |  |
| PsoOvis1B004289 | PSOVI05g00170 | 73% nAChR α-1 | 80% Dα1 | NI | 99% XM_027347514.1 | NI |
| PsoOvis1B004275 | PSOVI05g00180 | 64% nAChR α-1 | 69% Dα1 | NI | 84% XM_027347516.1 | NI |
| PsoOvis1B012767 | PSOVI52g00050 | 59% nAChR α-1 | 59% Dα2 | | 93% XM_027342746.1 | NI |
| PsoOvis1B000816 | PSOVI283g02900 | 71% nAChR α-3 | 50% Dα7 | | 78% XM_027341143.1 | NI |
| PsoOvis1B009263 | PSOVI69g00600 | 61% nAChR α-6 | 70% Dα7 | | 82% XM_027341343.1 | NI |
| PsoOvis1B001295 | PSOVI69g00580 | 68% nAChR α-5 | 73% Dα6 | | 62% XM_027341343.1 | NI |
| PsoOvis1B0146<br>36 | PSOVI283g043<br>00 | 79% nAChR<br>β-1 | 76% Dβ1 | | 99%<br>XM_027343735<br>.1 | NI |

Both *P. ovis* GluCls showed similar transcription patterns, with higher transcription levels in the larval stage (Figure 1, GluCl-44 218 reads per million and GluCl-280 192 reads per million) and, to a lesser degree, in protonymphs and adult males (83-119 reads per million). In the other two stages, the subunits had low levels of transcription, with between 10 and 33 reads per million. All other cysLGIC subunits had low transcription levels, between 1 and 50 reads per million. The phylogenetic analysis depicts the orthologues of the different cysLGIC subunits in Figure 2. DNA concentration after purification and Phusion^®^ high-fidelity PCR ranged from 33.6 ng/µl to 119.7 ng/µl. A detailed description of the samples can be found in Supplementary Table S3.

**Figure 1.**
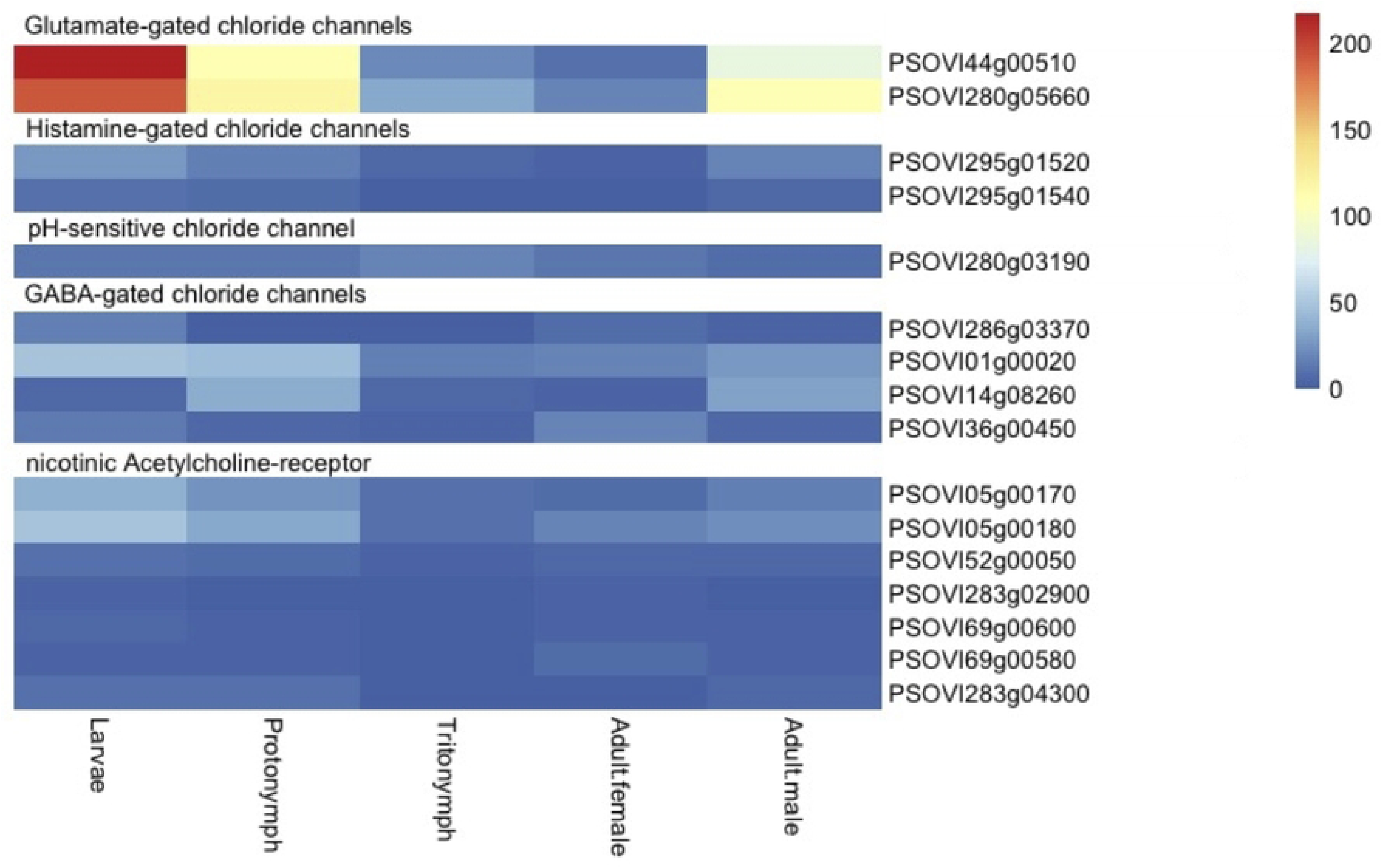
Heatmap of the transcription levels of the different *P. ovis* cysLGIC subunit encoding genes. Blue indicates low levels of gene transcription and red relative high levels of gene transcription.

**Figure 2.**
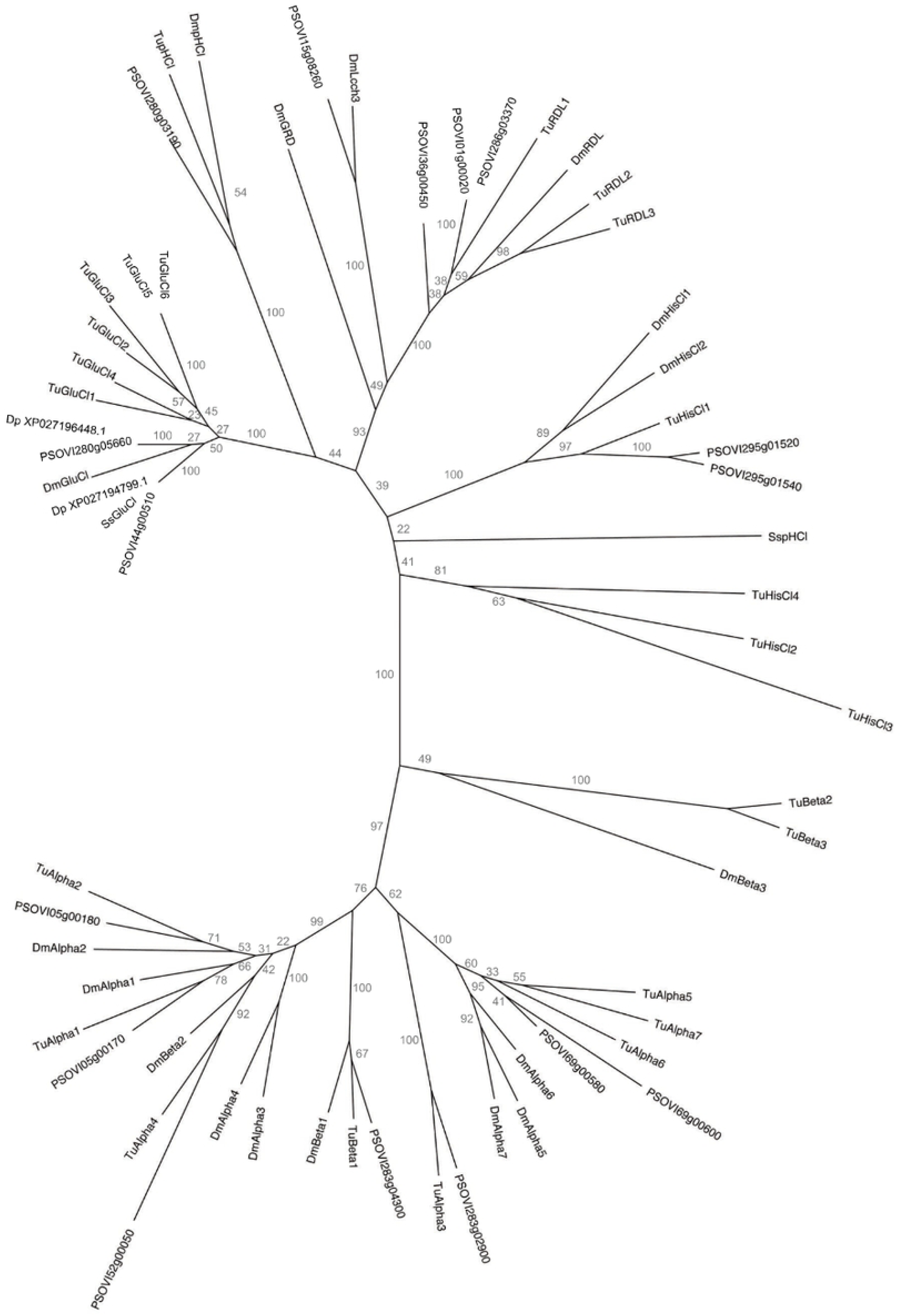
A maximum likelihood phylogenetic tree of cysLGIC subunits from *P. ovis* (PSOVI), *T. urticae* (Tu), *D. melanogaster* (Dm), *S. scabiei* (Ss) and *D. pteronyssinus* (XP_). The bootstrap values are given at the nodes.

#### 3.1.2. Deep amplicon sequencing

No variation was observed between the 179 bp-long sequences of the GluCl-44 gene and the GluCl-280 genes between the analysed mite populations. The amplified GluCl- 44 and -280 sequences were identical to both gene sequences previously submitted to OrcAE (Figure 3). Figure 4 illustrates how the location of these known mutations relates to the examined regions of GluCl-44 and GluCl-280.

**Figure 3.**
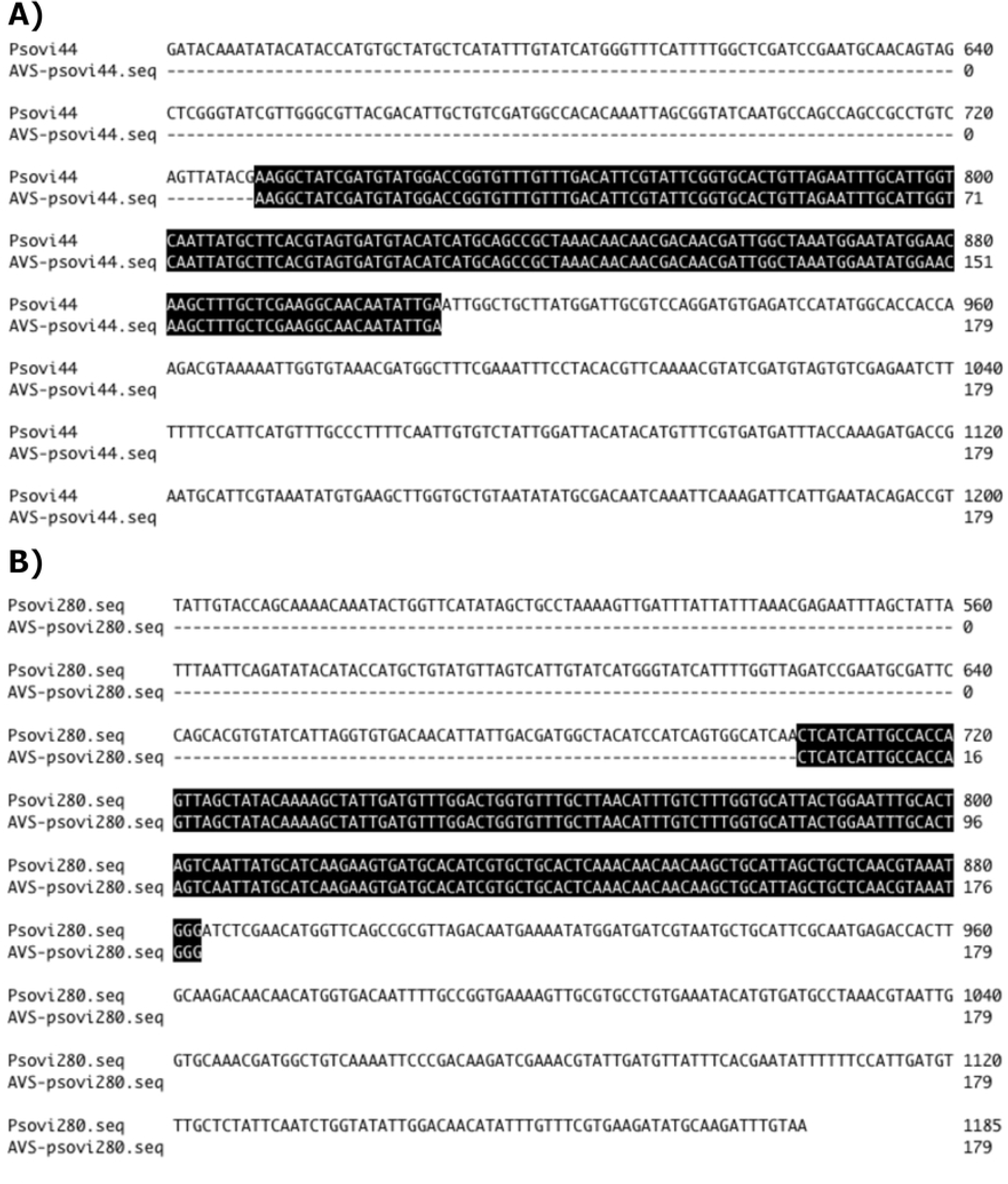
Alignments of the *P. ovis* GluCl-44 (A) and GluCl-280 (B) genes. Results of the deep amplicon sequencing ‘AVS-psoviXXX.seq’ and the OrcAE database ‘PsoviXXX.seq’.

**Figure 4.**
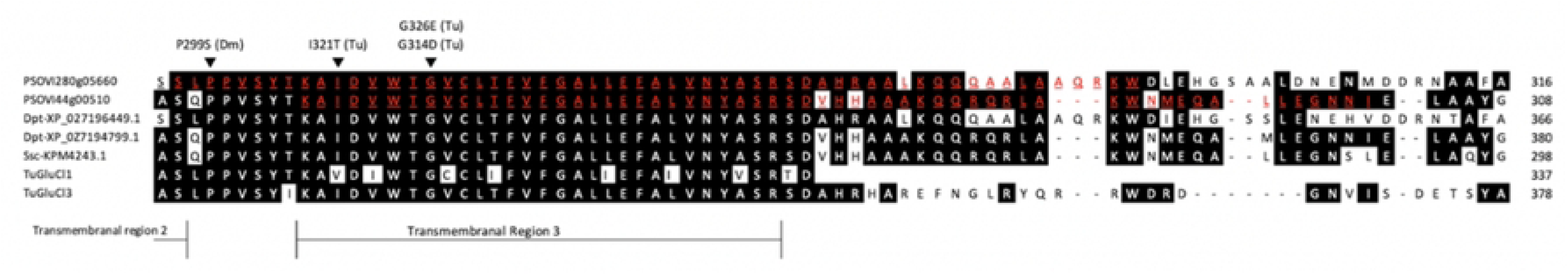
Alignments of glutamate-gated chloride channels from *P. ovis*, *D. melanogaster*, *S. scabiei* and *T. urticae*. The regions examined with deep amplicon sequencing are red underlined. Mutations from *D. melanogaster* (Dm) and *T. urticae* (Tm) associated with macrocyclic lactone resistance are given.

#### 3.2.1. Sample collection and RNA extraction

The post-treatment reduction in mite counts on the susceptible farm, Farm 2 from [13], was 97.4% with a 95%CI of [91.8%:99.9%]. The reduction post-treatment on the resistant farm, Farm 5 in [13], was 66.1% with a 95%CI of [55.1-75.6%]. Based on the criteria described in [13] the mite population on the first farm was classified as sensitive and the latter as resistant. RNA concentrations after extraction ranged from 102.6 ng/µl to 383.0 ng/µl and the RNA integrity number (RIN) was at least 8.3. A detailed description of the samples can be found in Supplementary Table S4.

#### 3.2.2 RNAseq analysis and identification of genes differentially expressed between *P. ovis* mite populations

RNA sequencing generated between 30-39 million reads per sample. The pseudo-alignment rates varied between 84-90%. Detailed information per sample is given in Supplementary Table S4. A total of 10,516 genes were tested in each comparison and the majority of genes were assigned an adjusted p-value with the RES_unexposed_ versus RES_exposed_ comparison having the least at 8,877 genes. DESeq2 results for each of the three comparisons between the three treatments are given in Supplementary Files S1-S3.

#### 3.3.1 Gene and isoform level differential expression

A Venn diagram, volcano plots and heatmap of differentially expressed genes with greater than log_2_-fold change of +1/-1 between the three pairwise contrasts is shown in Figure 5. Differential expression results implicated constitutively expressed detoxification genes in *P. ovis* ML resistance with 37 genes showing shared overexpression in RES_exposed_ and RES_unexposed_ versus SUS, 16 of which are doubled in expression versus SUS (> log-fold_2_ change of 1, Figure 5a). In both RES_exposed_ and RES_unexposed_ versus SUS comparisons a cytochrome P450 3A4-like gene (PsoOvis1B011549) was significantly highly overexpressed with (Table 3) absolute fold-changes of 4.9 and 3.3 respectively. This gene was also significantly over-expressed in the RES_exposed_ versus RES_unexposed_ comparison with 1.5-fold higher expression in RES_exposed_ replicates. Two UDP-glycosyltransferases (PsoOvis1B010992 and PsoOvis1B004414) were also highly overexpressed in RES_exposed_ and RES_unexposed_ versus SUS comparisons but not in the RES_exposed_ versus RES_unexposed_ comparison. An inositol oxygenase (PsoOvis1B000159) which may act in UGT detoxification is also constitutively overexpressed. These UGTs occur in tandem on chromosome 7 in the *P. ovis* sheep and cattle genomes. Several CYPs, UGTs and other detoxification genes including carboxylesterases, ATP-binding cassette and glutathione-S-transferase genes exhibited patterns of induced expression both up- and down-regulated in Res_exp_ (Table 3). Other potentially induced genes include a GRIN3B glutamate ionotropic receptor (PsoOvis1B009258). With respect to other possible resistance mechanisms two cuticular protein genes were overexpressed in RES_exposed_ versus SUS and RES_unexposed_ comparisons with 3.5-to-6-fold changes in abundance (Table 3, Supplementary Files S1-S3). A voltage-dependent T-type calcium channel subunit (PsoOvis1B011113) was downregulated in both RES_exposed_ and RES_unexposed_ versus SUS comparisons. An acetylcholinesterase-like gene (PsoOvis1B007848) was downregulated in RES_exposed_ versus RES_unexposed_ and SUS.

**Figure 5.**
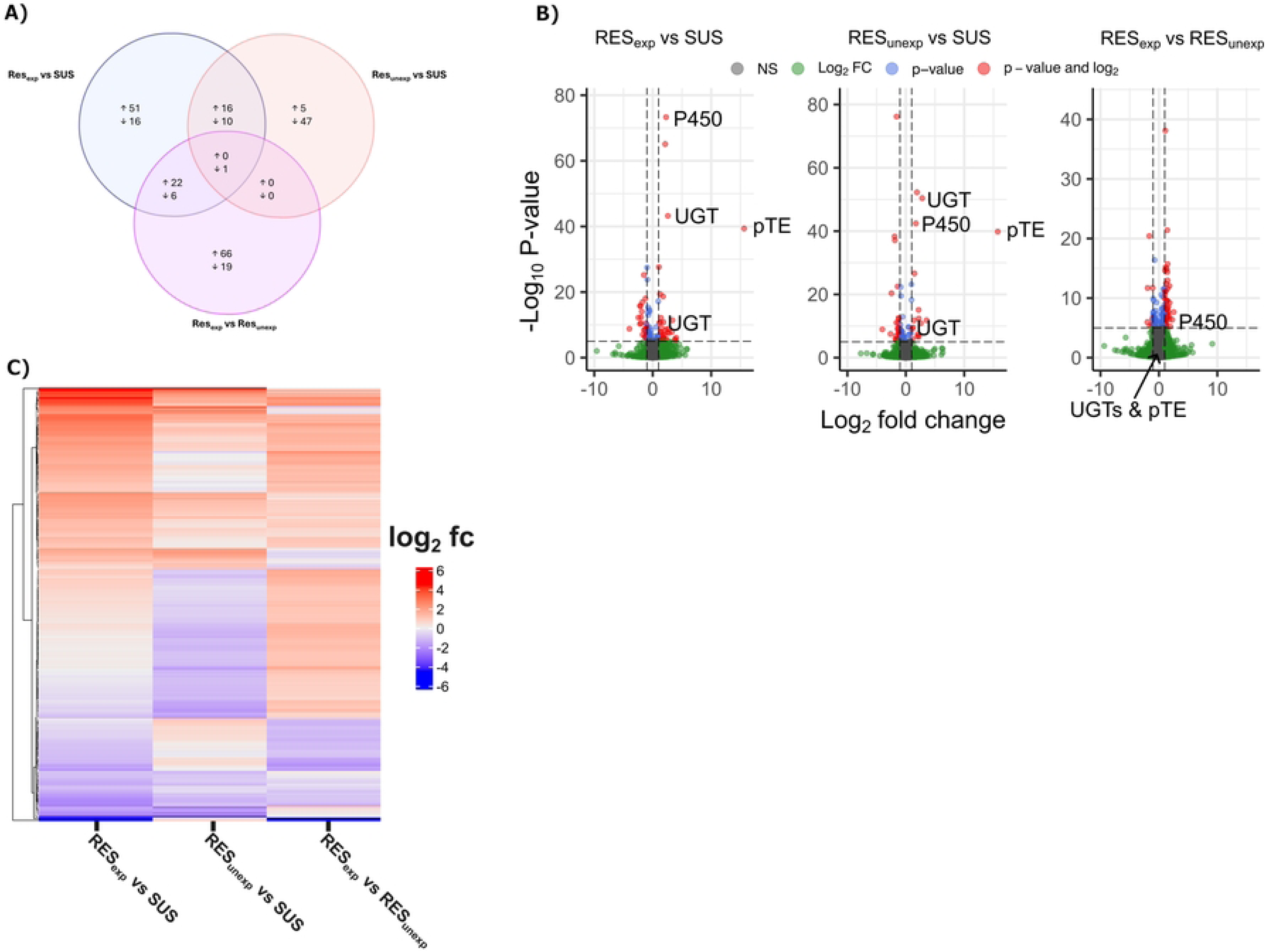
Summary of RNASeq results. A) Venn diagram showing concordantly up- (↑) and down-regulated (↓) genes between the and RES_exposed_ versus SUS, RES_unexposed_ versus SUS and RES_exposed_ versus RES_unexposed_ comparisons with greater than log-fold_2_ 1 difference. Up refers to genes more highly expressed in the first treatment labelled for each comparison. Shared differentially expressed genes between two comparisons are found in the overlapping circles. B) Volcano plots of differentially expressed genes for the three contrasts with key genes labelled: P450 = Cytochrome P450, UGT = *UDP*-glucuronosyltransferases, pTE = putative transposable element. C) Heatmap of log-fold change for each of the three contrasts undertaken for all genes included in (A). fc = fold-change.

**Table 3.** Overview of key differentially expressed genes for each comparison. R-exp = RES_exposed_, S = SUS, R-unexp = RES_unexposed_, fc = fold-change. A positive fold-change means upregulation in the first treatment (i.e. *1st vs 2nd*) of each comparison and a negative fold-change means upregulation in the second listed treatment.

| Gene Name | Predicted gene function | R-exp vs S FC | R-unexp vs S FC | R-exp vs R-unexp FC |
| --- | --- | --- | --- | --- |
| PsoOvis1B011549 | cytochrome P450 3A4-like | +4.9 | +3.3 | +1.5 |
| PsoOvis1B004414 | udp-glucuronosyl/udp-glucosyltransferase | +6.0 | +7.0 | -0.9 |
| PsoOvis1B011113 | voltage-dependent T-type calcium channel subunit alpha-1G | -0.5 | -0.6 | +0.9 |
| PsoOvis1B010992 | udp-glucuronosyl/udp-glucosyltransferase | +2.6 | +2.3 | +1.1 |
| PsoOvis1B002032 | Hypothetical transposable element | +∞ | +∞ | -0.9 |
| PsoOvis1B000159 | Inositol oxygenase | +2.8 | +4.0 | -0.7 |
| PsoOvis1B009258 | GRIN3B glutamate ionotropic receptor | +1.5 | -0.9 | +1.6 |
| PsoOvis1B007848 | Acetylcholinesterase | -0.6 | +1.2 | -0.5 |
| PsoOvis1B005563 | Argonaute protein | -0.7 | -0.9 | -0.8 |
| PsoOvis1B003877 | Argonaute protein | -0.7 | -0.8 | -0.9 |
| PsoOvis1B003033 | Cuticular protein CPR50 | +3.6 | -1.0 | +3.7 |
| PsoOvis1B004107 | Cuticle Protein | +3.6 | -0.6 | +5.2 |
| PsoOvis1B011188 | UDP-glucuronosyltransferase | +1.4 | +1.1 | +1.4 |
| PsoOvis1B009583 | Carboxylesterase | +1.4 | -0.9 | -0.5 |
| PsoOvis1B007144 | Carboxylesterase | +0.6 | -1.0 | -0.6 |
| PsoOvis1B006154 | Carboxylesterase | -0.5 | +0.9 | +1.8 |
| PsoOvis1B009247 | UDP-glucose 4-epimerase | +1.5 | +1.0 | +1.4 |
| PsoOvis1B008134 | glutathione S-transferase Mu 1 | +1.3 | +1.1 | +1.3 |
| PsoOvis1B002252 | glutathione S-transferase kappa 1 | +1.4 | +1.1 | +1.3 |
| PsoOvis1B007926 | cytochrome P450 4V2 | -0.5 | -0.7 | -0.7 |
| PsoOvis1B007062 | ABC transporter G family member 20 | +1.5 | +1.2 | +1.3 |
| PsoOvis1B000612 | ABC transporter G family member 20 | +1.3 | +1.1 | +1.1 |
| PsoOvis1B010281 | ABC transporter family C, ABCC3 | -0.6 | -0.9 | -0.7 |

Further genes of interest included two argonaute genes (PsoOvis1B005563 and PsoOvis1B003877) which were significantly downregulated in RES_exposed_ versus SUS. A gene likely to encode a retrotransposon, PsoOvis1B002032, had striking expression dynamics with high relative expression in RES_exposed_ and RES_unexposed_ and zero expression in SUS replicates with associated very low adjusted p-value. As is often the case for organisms such as *P. ovis*, with poorly characterised genomes, functional domain annotations were unavailable for many of the differentially expressed genes identified between both groups. Further investigation of their potential function could identify additional detoxification mechanisms.

At the isoform level results are concordant with the gene results with the exception of one of the five isoforms of the Gamma-Aminobutyric (GABA) transporter (PsoOvis1B011106, interpro accession: IPR000175, CDD accession: cd11496) which is down-regulated in both RES_exposed_ and RES_unexposed_ treatments versus SUS but is not significant in any gene level comparison (Supplementary Files S4-S6).

#### 3.3.2 Quantitative PCR results

The results of the qPCR confirmed the results of the RNA seq analysis with a positive, significant rmcorr estimated *r_rm_* value of 0.74 (95% confidence intervals 0.525-0.977, p<0.004) as can be seen in Figure 6A. The expression of inositol oxygenase (PsoOvis1B000159), a UDP-glycosyltransferase (PsoOvis1B010992) and the two cytochrome P-450s (PsoOvis1B011549 and PsoOvis1B005763) followed the same trend in qPCR as for the RNAseq data. Only for the other UDP-glycosyltransferase (PsoOvis1B004414) was the transcription pattern of the RNAseq data not confirmed by the qPCR, indeed removing this gene from the rmcorr increased the *r_rm_* value to 0.95 and narrowed the 95% CIs (Figure 6B, 0.729-0.992). This lack of concordance between qPCR and RNAseq data for PsoOvis1B004414 was unexpected, since the biggest differences in transcription after RNA seq were found for this gene.

**Figure 6.**
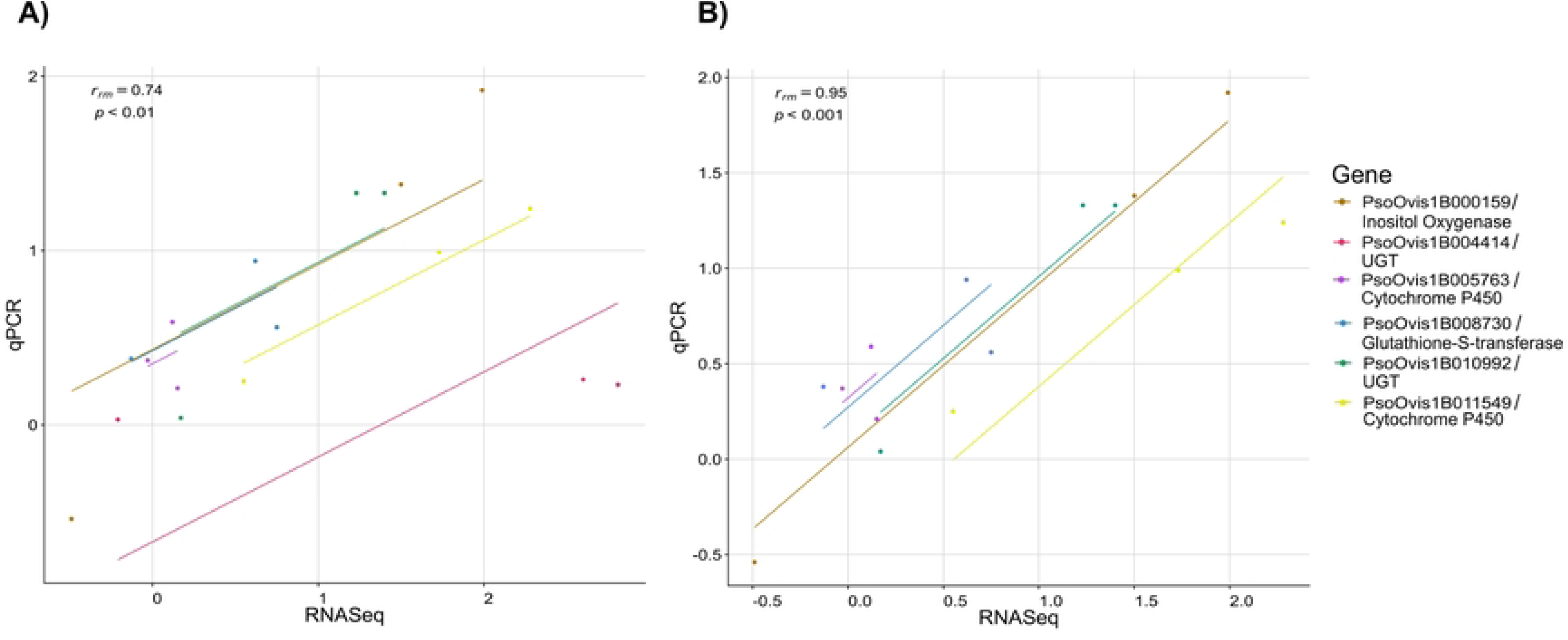
Rmcorr correlation analysis between log_2_ real-time quantitative PCR and RNASeq expression values for six selected genes. Each gene has three points representing the three experimental contrasts between RES_exposed_ RES_unexposed_ and SUS conditions, and a common trend line as fit by rmcorr to all genes. A) all six genes included correlation results: *r_rm_* = 0.74, 95% CI [0.525, 0.977], p = 0.004. B) Gene PsoOvis1B004414 removed from the analysis given the difference between RNASeq and qPCR values resulting in an increased positive correlation: r_rm_ = 0.95, 95% CI [0.729, 0.992], p < 0.001, as shown by the trend line. Raw data for this figure is given in Supplementary Table S13.

### 3.4 Population Genomics

The assembly span of sheep-derived *P. ovis* genome (62.7 Mb) and number of large scaffolds (10) are close to the expected size and number of chromosomes (Genome assembly metrics: Table 4). The less contiguous sheep-derived *P. ovis* assembly of [38] GCA_002943765.1, also has a span of 63 Mb and the Genomes on a Tree (GoaT, https://goat.genomehubs.org) is nine based on ancestral genomes. Although less contiguous, the cattle-derived *P. ovis* genome is of similar span (Table 4). Sequencing depths and alignment statistics are given in Supplementary Table S5. Of the 10,516 nuclear genes annotated in the sheep *P. ovis* genome (Table 5), 300 were not transferred to the *P. ovis* cattle genome assembly by liftoff. Of these, 206 genes lack informative annotations i.e. have “protein_coding”, “N/A”, hypothetical or uncharacterised annotation only. We capitalise the phenotype of the seven whole genome-sequence pooled template mite populations (Susceptible or Resistant) for clarity when referring specifically to these data. A further 58 encode transposable elements and 36 encode annotated genes. Pairwise genome-wide genetic distances, F_st_ and PCA between the Susceptible and Resistant populations indicate separation between the two phenotypes and high-similarity among the Resistant populations (Figure 7).

**Figure 7.**
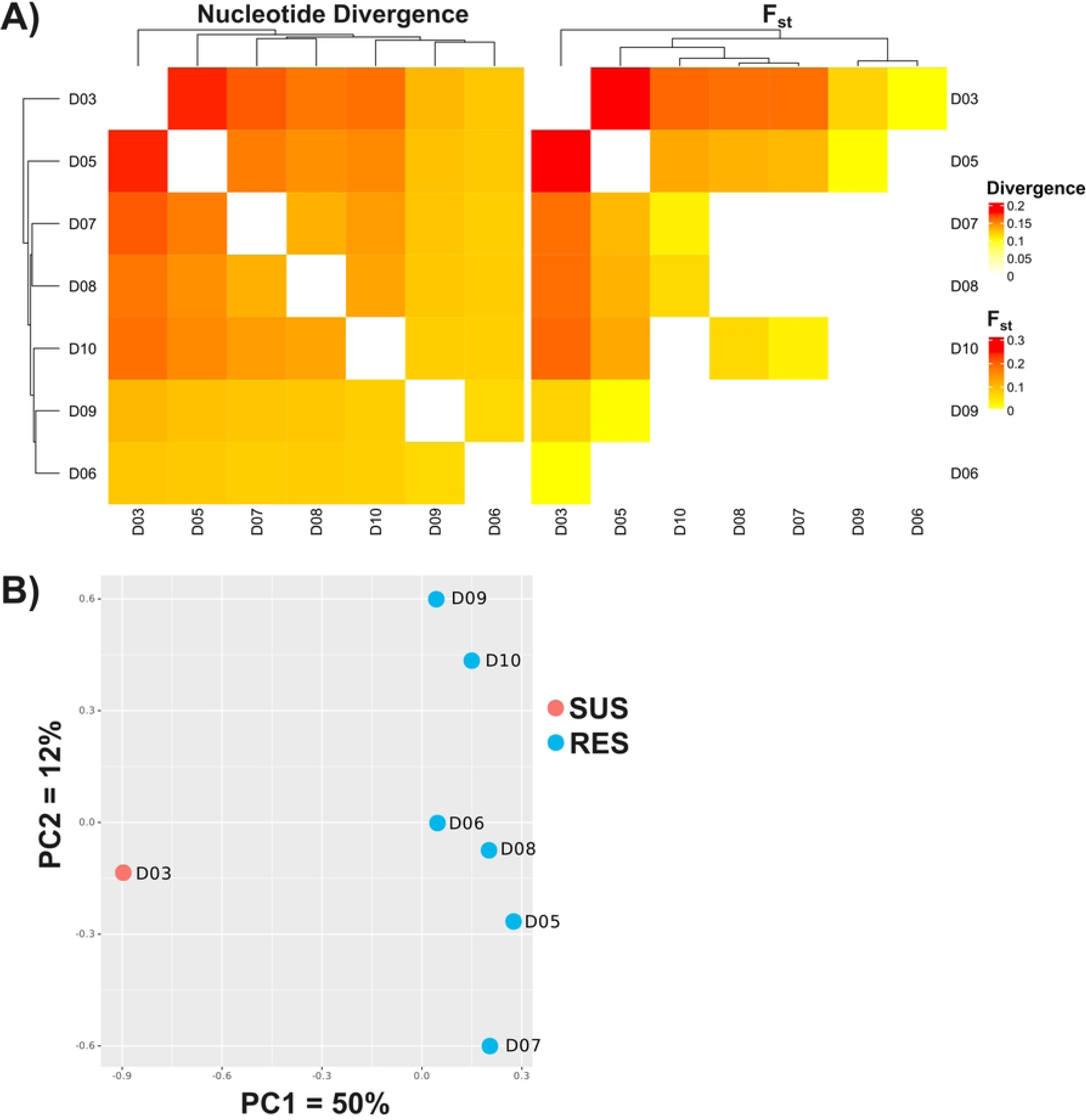
Relatedness of genome re-sequenced *P. ovis* populations. **A)** Heatmap of global pairwise nucleotide divergence and Fst between all whole genome sequenced pooled template populations included here. D03 = Susceptible; D05, D06, D07, D08, D09 & D10 = Resistant. **B)** Principal components analysis for the first two components for distance between populations coloured by phenotype and labelled by population. X-axis explains 50% of variance and discriminates between the Susceptible population and Resistant populations.

**Table 4.**
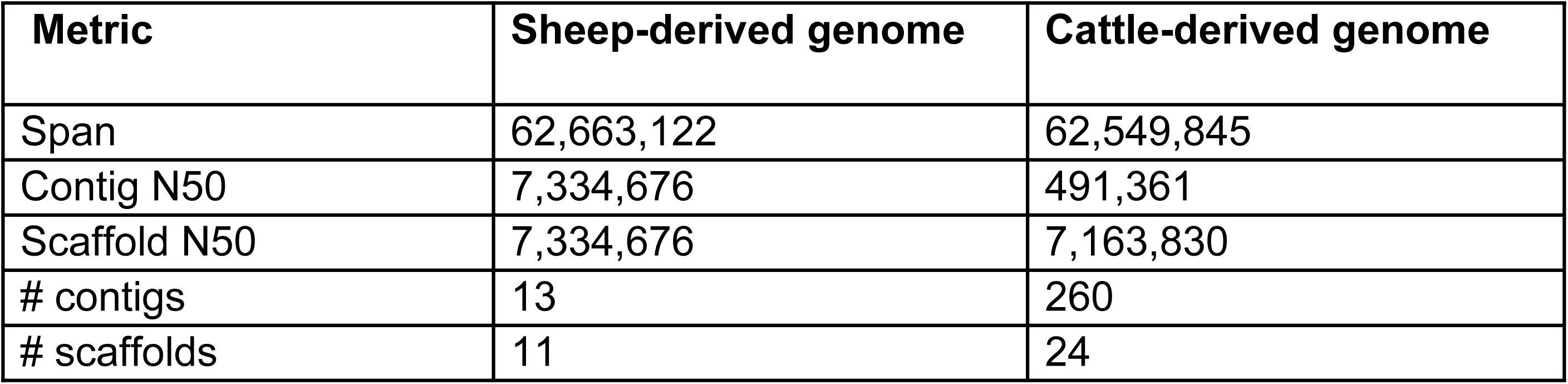
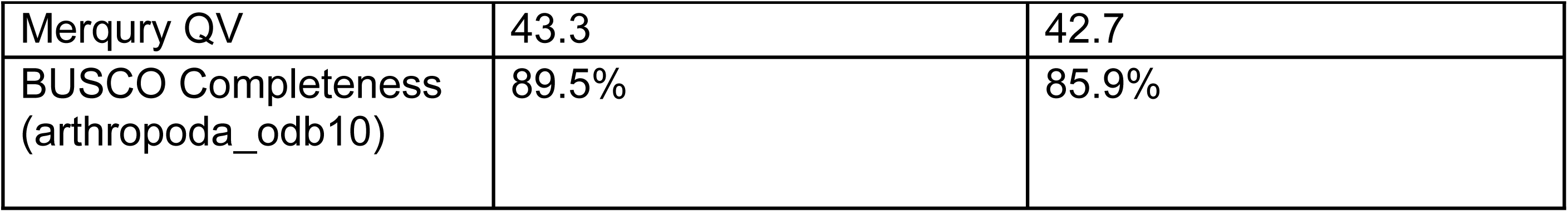
Genome assembly metrics for sheep- and cattle derived *Psoroptes ovis* assemblies. Merqury QV = Merqury consensus quality value [75]. BUSCO = Benchmarking Universal Single-Copy Orthologs [76]

**Table 5.**
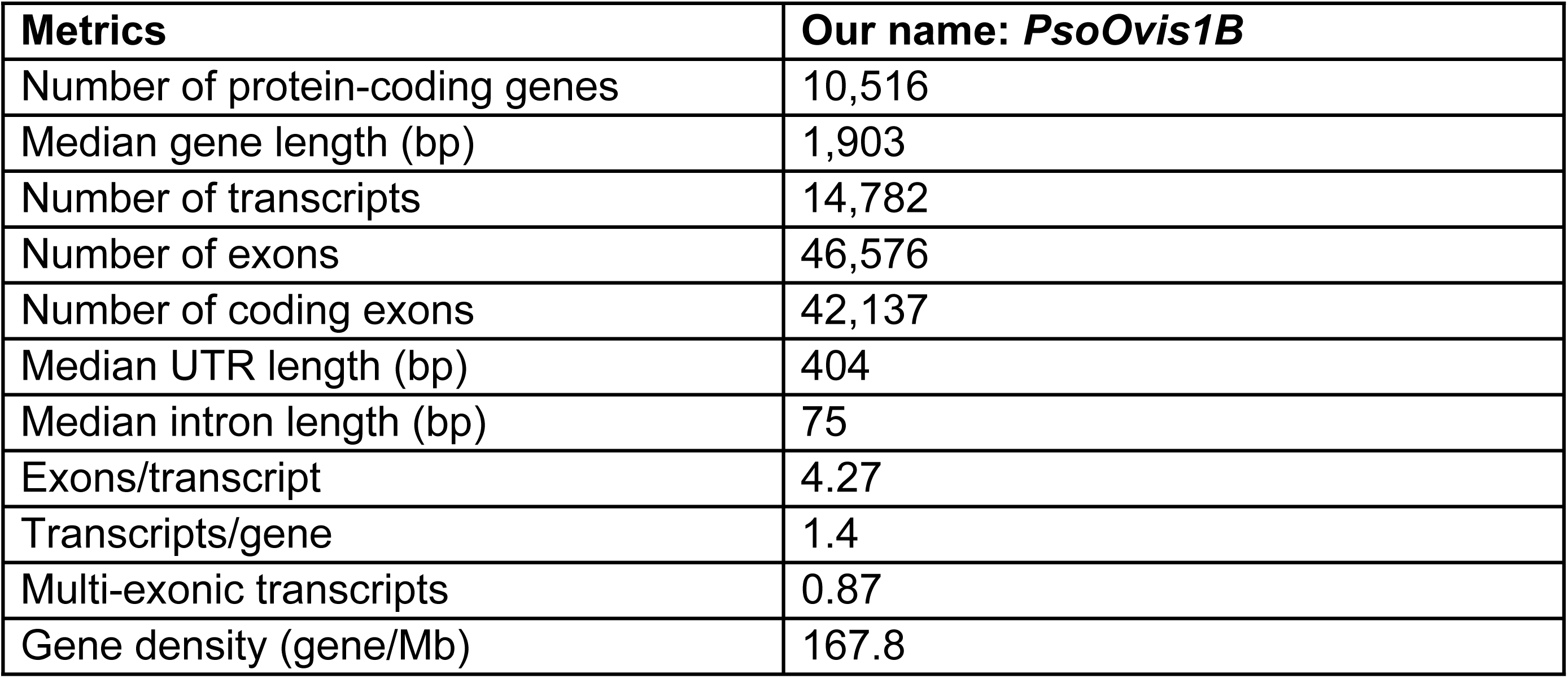
Genome annotation metrics for *P. ovis* used in RNASeq and population genomic analyses.

| Metrics | Our name: *PsoOvis1B* |
| --- | --- |
| Number of protein-coding genes | 10,516 |
| Median gene length (bp) | 1,903 |
| Number of transcripts | 14,782 |
| Number of exons | 46,576 |
| Number of coding exons | 42,137 |
| Median UTR length (bp) | 404 |
| Median intron length (bp) | 75 |
| Exons/transcript | 4.27 |
| Transcripts/gene | 1.4 |
| Multi-exonic transcripts | 0.87 |
| Gene density (gene/Mb) | 167.8 |

#### 3.4.1 Gene copy number variation

Elevated coverage for each population was identified at two loci across the *P. ovis* cattle genome through comparison of average gene coverages and visual inspection of read data in IGV. On Chromosome 2 the Cytochrome P450 3A4-like gene PsoOvis1B011549 with high-expression and a neighbouring short-chain dehydrogenase gene PsoOvis1B003189 have elevated coverage in all Resistant populations but not the Susceptible population where coverage is not different from the genome-wide median (Figure 8). The boundaries of the 3,347 bp elevated coverage region are clearly defined at positions 3,606,198 to 3,609,545 in the cattle *P. ovis* genome (Figure 8). For each of the six populations coverage is even across the elevated region including introns present in both PsoOvis1B011549 and PsoOvis1B003189 genes. Coverage varies widely as a multiple of the median across Resistant populations indicating possible variable copy numbers across sampled populations (Figure 8). Gene PsoOvis1B003189 was also over-expressed in both RES treatments versus SUS but not in RES_exposed_ versus RES_unexposed_. On Chromosome 7, the tandemly located UGTs PsoOvis1B010992 and PsoOvis1B004414 exhibit even more extreme coverage dynamics (Figure 8). In contrast to the high-copy number CYP gene, the Susceptible population also exhibits increased copy number of the two UGT genes but without associated gene over-expression. Average coverage/median coverage is below 50 in these genes in the Susceptible population but above this level in all Resistant populations. The alignment profile of the elevated region also differs between Susceptible and other populations (Figure 8).

**Figure 8.**
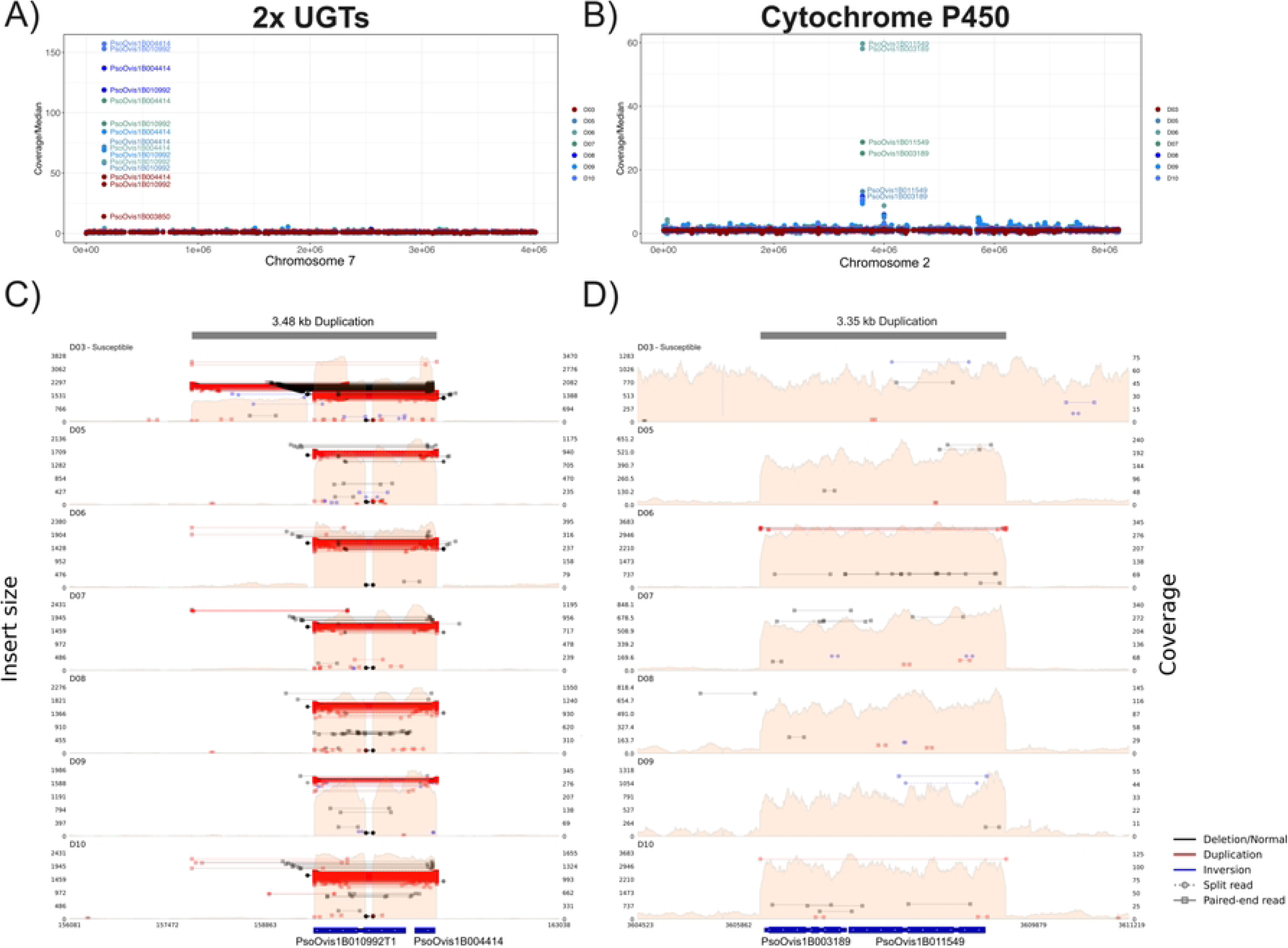
Copy number variation of the over-expressed UGT (PsoOvis1B004414 and PsoOvis1B010992) and cytochrome P450 gene (PsoOvis1B011549). A) & B) Coverage of each gene divided by median genome-wide coverages per population sequenced for the two chromosomes encoding the two UGT and cytochrome P450 genes. Resistant populations are coloured blue and green and Susceptible in red, for B) gene labels were removed for several Resistant populations as they were overlapping. C) & D) Read alignment patterns across these loci showing insert size of paired end reads and overall read coverage (pink regions) with coding region locations for UGTs and the cytochrome P450 plotted beneath the X-axis.

Other loci with elevated coverage in Resistant versus the Susceptible population include PsoOvis1B001111 also on Chromosome 2, a GTP-ase activating protein, although this gene is not differentially expressed in any comparison. For all populations, elevated average coverage of this gene is explained by a 19 bp spike in coverage against background genome coverage, possibly resulting from transposable element activity. A plexin A gene (PsoOvis1B001325) on Chromosome 2 is elevated in all populations but more so in Resistant than Susceptible in *P. ovis* cattle genome aligned data but, surprisingly, not the *P. ovis* sheep genome (Supplementary Tables S6 and S7). This gene was not differentially expressed in any comparison. Two genes (PsoOvis1B007656 & PsoOvis1B000689) at the 3’ end of Chromosome 4 are also elevated in coverage, one of which encodes for a reverse transcriptase. Finally, a gene on Chromosome 1 (PsoOvis1B004923) and Chromosome 10 (PsoOvis1B007634) were also elevated versus background coverage. Another gene with a distinct coverage profile is Skeletor (PsoOvis1B008827) which is elevated in alignments against the sheep but not the cattle *P. ovis* genome.

We used nanopore data to explore possible cis copy number variation through the occurrence of multiple copies of a gene on the same long read for both sheep and cattle datasets. For sheep nanopore data, CYP gene PsoOvis1B011549 only occurs once in every read whereas 8 reads had evidence for more than one copy of PsoOvis1B011549 and PsoOvis1B003189 in cattle data (Supplementary Table S8). By contrast many reads had non-overlapping consecutive matches to the two UGT genes in the cattle data. The longest nanopore read (eb6fdcaf-bc66-4353-a602-9c8fa9087462) at >70Kb encoded 40 copies of PsoOvis1B010992 with matches to three further genes including PsoOvis1B000689 (Figure 9), 35 of the genes occurred consecutively along the read (Supplementary Table S9). The read (17f558b9-9b0f-4336-94f5-3de162674cb6) with most copies of UGT PsoOvis1B004414 encodes 20 copies of the gene alongside 20 copies of PsoOvis1B010992 and PsoOvis1B014440. In the sheep nanopore data the most copies of either UGT encoded by a single read was three copies (Supplementary Table S10).

**Figure 9.**
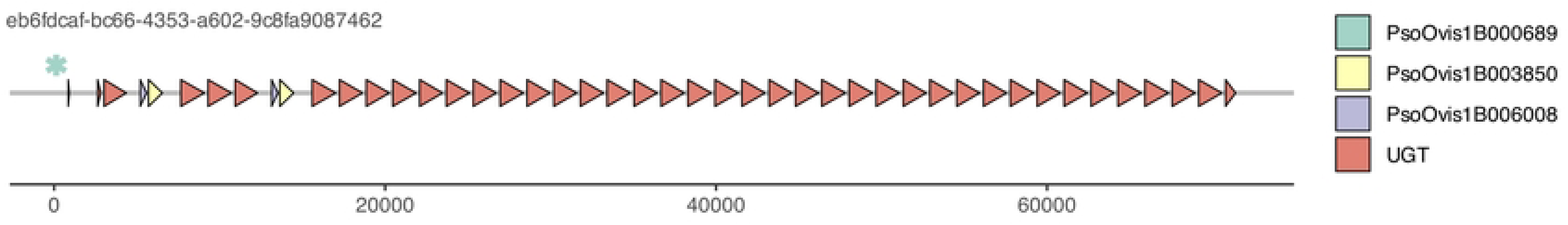
Multiple copies of a UGT gene along an individual long-read sequence for the read with the most individual copies. A coloured asterisk is given for PsoOvis1B00689 as it is too short to colour the gene arrow.

Read alignments around PsoOvis1B011549 were inspected in IGV to identify the mode of copy number increase in the Resistant versus the Susceptible populations. The increase in coverage was clearly delineated in Resistant populations with no change versus surrounding sequence in the Susceptible (Figure 8). Discordant read mappings at the start and end of the sequence with elevated coverage were used to identify potential insertion positions in other regions of the *P. ovis* genome for both cattle and sheep assemblies. Five sites were identified with elevated coverage of ∼250bp length in all Resistant populations but not the Susceptible population. Localised increases in read number concordant with putative insertion sites and confirmed by inspection in IGV (Supplementary Figure S1). Two of the potential insertion sites flank either side of a gene, PsoOvis1B003901, on chromosome 4. One further CYP gene on Chromosome 5, PsoOvis1B000586, had elevated coverage in one Resistant population (D06) in *P. ovis* sheep genome data only, as this gene was not placed on the *P. ovis* cattle genome assembly (Supplementary Tables S6 & S7). Although significantly differentially expressed in two comparisons (RES_exposed_ versus RES_unexposed_ and RES_unexposed_ versus SUS) it is not a gene with high absolute expression. Furthermore, this gene and two 3’ loci (PsoOvis1B006329 & PsoOvis1B014010) were not placed by the liftoff annotation into the cattle *P. ovis* assembly. Few other copy number variants correlate with resistance phenotype, including a deletion of 162bp not found in the Susceptible population in an intergenic region of Chromosome 5 (position 1,943,240 cattle genome).

#### 3.4.2 Genome wide F_st_ scans

Many peaks of differentiation were evident in pairwise F_st_ scans between Resistant (D05-D10) and Susceptible (D03) populations (Figure 10). This was also true in intra-Resistant population comparisons. Chromosomes 1 and 2 as the largest chromosomes contain the most signals of differentiation. Given the multiple pairwise comparisons, we decided to restrict the analyses to genes with average Fst differences of 0.2 or greater between Susceptible versus Resistant pairwise comparisons and Resistant versus Resistant comparisons. In this way we were able to focus on genes that had a consistent signal of differentiation between Susceptible and Resistant comparisons (x6) while avoiding differentiation shared with Resistant-only population pairwise comparisons (x15).

**Figure 10.**
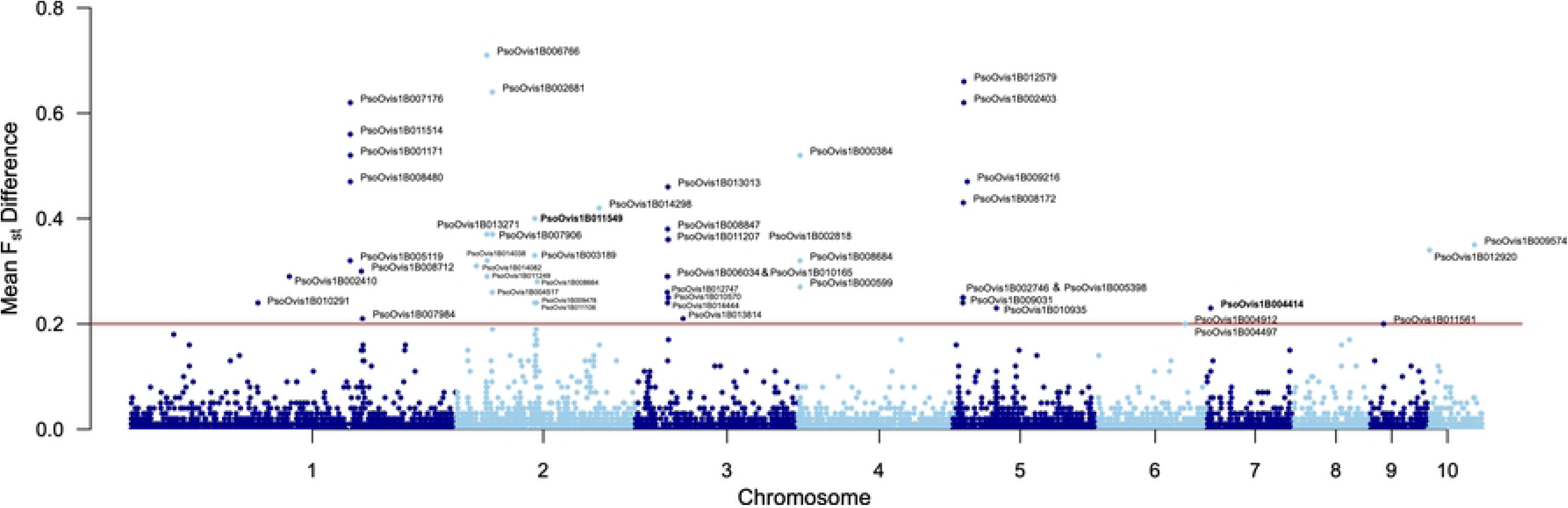
Per gene absolute mean difference in F_st_ (|F_st_|) between Resistant versus Susceptible comparisons and Resistant versus Resistant comparisons. Chromosomes are numbered along the X-axis with a change in colour indicating a break between chromosomes. A cut-off of 0.2 was applied shown by the horizontal red line, genes above this are labelled with font size changed to avoid overlap.

Only 48 loci passed the threshold for consideration (Figure 10, Supplementary Table S11) using the cattle genome aligned dataset. This included the cytochrome P450, PsoOvis1B011549, which has an average of F_st_ of 0.4 across the six D0X versus D03 comparisons versus 0.0 for intra-Resistant comparisons giving an absolute difference (|F_st_|) of 0.4. One of the two UGTs, PsoOvis1B004414, was also in the 48 genes at |F_st_| 0.23 versus |F_st_| 0.06 for PsoOvis1B010992. The Gamma-Aminobutyric (GABA) transporter (PsoOvis1B011106) which exhibits differential transcript usage also had high |F_st_| at 0.24. Only four genes overlap with differentially expressed genes for RES_exposed_ and RES_unexposed_ versus SUS comparisons. These included the cytochrome P450 PsoOvis1B011549 and adjacent amplified gene PsoOvis1B003189, the UGT PsoOvis1B004414 and PsoOvis1B013814 on Chromosome 3 which is highly overexpressed in RES_exposed_ and RES_unexposed_ versus SUS at 3.5 and 2.6-fold greater expression respectively. This gene is not annotated and does not encode a diagnostic protein domain (InterProScan search) and is most similar to clumping factor B-like in *Dermatophagoides farinae* by protein blast search, albeit at a relatively weak e-value 1 x 10^-8^.

Extended regions of |F_st_| > 0.2 occurs on Chromosome 2 at ∼1 mb and Chromosome 3 at ∼1.45 mb (Figure 10). Genes in these regions weren’t correlated to resistance phenotype by gene expression or copy number variation analyses. Some genes with high |F_st_| encode nonsynonymous mutations with inverse frequencies between the Susceptible and Resistant populations (Supplementary Table S12).

#### 3.4.3 Variants associated with ML-resistance candidate loci

No non-synonymous mutations were detected in the high-copy number, over-expressed CYP genes in any populations. Three synonymous mutations occur at close to 100% frequency in the six Resistant populations which are not present (0% frequency) in the Susceptible population (Supplementary Table S12). A 5’ upstream variant in the amplified DNA region is fixed in the Susceptible population but not present in any other populations. The neighbouring short-chain dehydrogenase gene PsoOvis1B003189 that shares elevated coverage with PsoOvis1B011549 encodes a premature stop codon at amino acid 391 of 798. This stop codon is at 100% frequency in all Resistant populations but does not occur (0% frequency) in the Susceptible population. The UGT gene PsoOvis1B010992 encodes two frameshift mutations within three base pairs of each other. These mutations occur at intermediate levels in the Susceptible population at 34% and 37% respectively whilst occurring at low prevalence (<5%) in all other populations (Supplementary Table S12). As they involve a two base pair insertion and deletion respectively it is not clear if a frameshift results in the final protein. Three missense mutations follow a similar pattern of prevalence with intermediate levels in the Susceptible population and low in the six Resistant populations.

The Gamma-Aminobutyric (GABA) transporter (PsoOvis1B011106) encodes many mutations between the Susceptible and Resistant populations; however, none result in nonsynonymous changes. We did not identify any non-synonymous mutations in the whole genome re-sequencing data.

## 4. Discussion

### 4.1. Target site variation

The observed number of cysLGICs is less than in other arthropod species such as *T. urticae* (n=29), *D. melanogaster* (n=23), the wasp *Nasonia vitripennis* (n=26) and the honeybee *Apis mellifera* (n=21) [21,77,78]. Two genes encoding for a GluCl-subunit were identified in *P. ovis*. This varies from the other examined species, as *T. urticae* has 6 genes encoding for GluCl subunits, *D. melanogaster* 1, *S. scabiei* 3 and *D. pteronyssinus* 4 [21,79]. Two HisCl subunits were also found in *D. melanogaster*, the honeybee *A. mellifera* and the parasitoid wasp *Nasonia vitripennis.* Only the two spotted spider mite, *T. urticae*, is known to have 4 different subunits [21,77,78,80]. Less variation in subunit numbers was observed with the pH-Cl. *Psoroptes ovis* has just one subunit, which is the same as that observed in *S. scabiei, D. melanogaster, A. mellifera, N. vitripennis* and *T. urticae* [21,25]. Previous functional expression of the pH-Cl from *S. scabiei,* showed activation of the channel after binding with ivermectin [25].

Four genes that encode for GABA-Cl subunits were characterised (Table 2). Three of which had high similarities with the RDL-subunit from *T. urticae* and *D. melanogaster* and the GABA-Cl β subunit from *D. pteronyssinus*. Mutations in the RDL-subunit cause dieldrin resistance in *D. melanogaster* [81]. The final gene, PsoOvis1B001590, shared high similarity with the Lcch3-subunit from *D. melanogaster* and the GABA-Cl α subunit from *D. pteronyssinus*. The Lcch3-subunit alignment shared a high similarity with the vertebrate GABA receptor β subunit [81,82].

Furthermore, 7 genes were characterised as nAchR subunit encoding genes. The only one classified as a β subunit being PsoOvis1B014636. nAchR β subunits lack 2 adjacent cysteine residues in loop C, that are present in the α subunits and other cysLGIC. The others more closely resembled α subunits. PsoOvis1B000816 has a high homology with nAChR □-3 of *T. urticae.* This subunit had an atypical FxCC amino acid sequence in the conserved C loop, instead of the highly conserved YxCC. This unusual amino acid sequence results in a decrease in acetylcholine affinity [21,23,83]. It occurs twice that two nAchR subunit encoding genes are in close proximity to each other in the genome and have a high homology.

Compared to the observed reads of *T. urticae* in OrcAE, the transcription levels of the cysLGICs in *P. ovis* are within the same range and magnitude, as can be seen in Supplementary Table S3. The low transcription of the non GluCl cysLGIC is not unexpected, as cysLGIC are present in a limited number of cell types, e.g. HisCl most likely in light-sensitive sensory neurons [21,22].

The presence of two GluCl subunits and their similar transcriptional patterns are possibly indicative of a past duplication event. The close proximity and high homology between some of the cysLGICs are indicative of gene duplication. Within this ion channel superfamily, a duplication event was found in the GluCls, HisCls and two in the nAchRs (PsoOvis1B004289 and -00180; PsoOvis1B009263 and -00580). The absence of any variation observed after deep amplicon sequencing can be expected from the ovine mite population, as this is the same mite strain that was used to build the OrcAE database. Despite this, the absence of any variation within the population is remarkable. Moreover, the sequences of the 8 cattle mite populations were also identical to the ovine mite populations. The amplicon of GluCl-44 covered parts of TM2 and TM3 and GluCl-280 covered TM3, regions known to contain mutations correlated with ML resistance in multiple arthropods and nematodes [27]. The absence of variability in these regions makes it less likely that mutations in GluCls are involved in ML resistance. In other arthropod species, mutations in GluCl genes have been associated with ML resistance in *D. melanogaster* and *T. urticae* [21,84,85].

### 4.2. Pharmacokinetic changes

Through a combination of gene expression and population genomic data our results show a strong correlation between macrocyclic resistance in cattle-derived *Psoroptes ovis* and combined overexpression and increased copy number of a cytochrome P450 and two tandemly located UDP-glycosyltransferase genes. Although other resistance-associated results emerge from our data, these detoxification genes emerged from analysis we undertook and are responsible for resistance to multiple families of pesticide in many arthropod species, including other mites species [35,86–92].

#### 4.2.1 Overexpression of metabolic resistance genes

The most prominent genes emerging from the RNAseq experiment were two UDP-glycosyltransferase (UGTs) and a CYP gene which were constitutively up-regulated in all six RES_exposed_ and RES_unexposed_ mite replicates. Although recognised as important components of metabolic resistance to pesticides in arthropods the role of UGTs is less well-characterised than CYPs [92–95]. Our study is one of the first descriptions of UDP-glycosyltransferase expression in an arthropod species of veterinary importance, as they are absent in some well-known non-insect arthropods, such as the *Ixodes* and *Rhipicephalus* ticks, the honeybee mite, *Varroa destructor*, and the predatory mite, *Metaseiulus occidentalis* [93,94]. Indeed, UGTs were lost early in chelicerate evolutionary history, but were re-acquired from bacteria in mites [94,96]. In the spider mites, *T. urticae* and *Tetranychus cinnabarinus,* up-regulation of a UDP-glycosyltransferase was linked to resistance to the ML abamectin [92,94]. Multiple recombinant UDP-glycosyltransferases from *T. urticae*, expressed in *E. coli*, were capable of metabolising abamectin and milbemectin *in vitro.* Furthermore, inhibition of a UGT gene with RNAi resulted in increased mortality in abamectin resistant strains of *T. cinnabarinus* [92,95,97]. The constitutive up-regulation of an inositol oxygenase orthologue (PsoOvis1B000159) also indicates a possible role of UDP-glycosyltransferase in the detoxification of macrocyclic lactones. This enzyme catalyses the production of D-glucuronate from inositol. D-glucuronate is involved in mammalian Phase II detoxification processes and could potentially have a similar role in invertebrates [98]. Snoeck et al. (2019) [95] observed the use of UDP-glucose as a main donor in *T. urticae*, but UDP-glucuronic acid was also utilised by certain UDP-glycosyltransferases. To fully understand the role of this gene in *P. ovis* further studies are required to provide a functional characterisation. Results from qPCR broadly support the RNASeq results with only PsoOvis1B004414 having differing expression dynamics across the three treatments. In SUS mites, this gene was also expressed highly, to a similar extent to the RES_exposed_ and RES_unexposed_ mites.

#### 4.2.2. Combined overexpression of detoxification genes

Combined over-expression of Class I (CYPs) and Class II (UGTs) detoxification genes observed here occurs in other arthropod pests and vectors resistant to a variety of pesticides. Examples include the colorado potato beetle, *Leptinotarsa decemlineata*, in response to the neonicotinoid imidacloprid [87], the green peach aphid, *Myzus persicae*, against sulfoxaflor [90] and the beet armyworm, *Spodoptera exigua*, in response to multiple insecticides including abamectin [86]. Interestingly, for *L. decemlineata* silencing the CYP and UGT genes through RNAi did not improve the mortality of imidacloprid versus RNAi of each gene individually. UGTs specifically have been implicated in resistance to organophosphates in the house fly *Musca domestica* [99]. This is concerning as plunge dipping in the organophosphate, diazinon, is heavily relied upon for the effective control of sheep scab. This method of treatment has also been shown to be highly efficacious against ML-resistant sheep-derived *P. ovis* mites (S. Burgess, Personal Communication).

#### 4.2.3. Other genes of interest

We observed significant, but modest, differences in expression of further detoxification pathway genes including glutathione S-transferases, ATP binding cassette, carboxylesterase and further CYP genes between the SUS, RES_exposed_, and RES_unexposed_ *P. ovis* isolates (Supplementary Files S1-S3). In *S. scabiei* and *T. urticae,* glutathione S-transferases of the µ- and ∂-classes have been linked to ML resistance, while a glutathione S-transferase of the ∂-class was capable of conjugation of abamectin [100,101]. The presence of environmentally induced genes offers phenotypic flexibility that makes it possible to adapt to new environmental challenges [102,103]. However, for the development of a potential diagnostic test for resistance, constitutively over-expressed genes would result in less-labour intensive diagnostics. As these have the possibility to be detected in acaricide RES_unexposed_ populations, while treatment induced over-expressed genes need acaricide exposure for their detection.

#### 4.2.4. No differential expression of GluCls

Finally, no cys-loop ligand gated ion channel component genes were found among the significant genes, nor were resistance-associated mutations discovered in the amplicon regions examined or whole genome re-sequencing data. This superfamily of arthropod ion-channels is regarded as the main target site of the MLs [31,104]. Although not of the same family of ion channels, in addition to differential expression, functional expression in *Xenopus laevis* showed that mutations can lower the susceptibility of GluCls for ivermectin [24]. The higher transcription of GluCls in the adult males and protonymphs (Figure 1) could be explained through the increased motility of these stages [39], as the main functions of GluCls in invertebrate nervous systems are control and modulation of locomotion, the regulation of feeding, and the mediation of sensory inputs [22]. However, an explanation for the higher levels in the larval stage is lacking. In lice, GluCls appear to play a role in feeding behaviour [105]. However, the low expression of GluCls in tritonymphs, a major feeding stage, is not in line with this finding [39].

We did however, observe isoform level differential expression and elevated F_st_ of one isoform of a GABA transporter gene (PsoOvis1B011106) without associated differential gene expression of this highly expressed gene. This suggests a change in the proportion of isoforms of this gene versus whole gene expression in RES samples. GABA transporter genes belong to three families and help regulate the concentration of extracellular GABA. The ML abamectin is believed to activate GABA transporters which results in excessive GABA release and ultimately death of the target organism [106,107]. The role of differential transcript usage in acaricide, and wider pesticide resistance in arthropods, is little understood currently but results such as ours could form a basis for functional investigations. There was evidence for induced expression of an NMDA-selective glutamate receptor (PsoOvis1B009258) gene.

### 4.3. Increased copy number in UGT and CYP genes

The increase in gene copy number observed for the two over-expressed UGT genes was observed in all whole genome sequenced populations of cattle-derived *P. ovis*, including the Susceptible. Although our data are not conclusive, the pattern of UGT genes in long reads from the *P. ovis* sheep and cattle genome assembly dataset indicates this increase in UGT genes may differentiate cattle mites from sheep. It is not clear whether repeated tandem duplications of these genes have occurred *in-situ* or if blocks encoding multiple gene copies have been dispersed around the genome through a process of segmental duplication. Future work involving targeted long-read sequencing and/or fluorescence *in-situ* hybridisation (FISH) of target UGTs would help to answer this question. The gene coverage plots indicate fewer copies in the Susceptible population than Resistant populations meaning copy number could be dynamic for UGTs, but this metric may not be sensitive to sequencing depth variability and other sources of bias and therefore requires confirmation. There are no fixed variants that distinguish Susceptible from Resistant populations, although intermediate frequencies of variants present in the Susceptible population suggest a distinct haplotype of these genes is segregating in that population. Unfortunately, the pooled template sequencing we undertook can only provide allele frequency data as haplotype information is lost, hence we cannot confirm if a susceptible-only haplotype is present which would explain these intermediate allele frequencies. The lack of overexpression of UGTs in the susceptible (SUS) RNASeq replicates suggests differences in gene regulation occurs at these loci between Susceptible and Resistant mites. However, in the qPCR results PsoOvis1B004414 was overexpressed in Susceptible mites to a similar extent as Resistant mites. This could result from a technical issue, for example lack of specificity in the qPCR primers or sequence differences at this locus between sheep and cattle-derived *P. ovis* genomes given that we have used the sheep genome annotations in this study. In addition to copy number changes, we observed distinct polymorphisms between Susceptible and Resistant populations including potential frame-shift mutations which may contribute to the observed differences in expression through effects on gene regulation.

#### 4.3.1 Gene amplification in subtelomeric regions

Multiple rounds of gene amplification as we observed here, also explains resistance to the herbicide glyphosate in the parasitic weed, *Amaranthus palmeri* [108,109]. The 5-enolpyruvylshikimate-3-phosphate synthase (EPSPS) locus is amplified 100-fold as part of a 400 kb long cassette of the EPSPS gene and 58 other genes. This EPSPS encoding cassette of genes occurs as extrachromosomal circular DNA and putatively originated from the activity of a transposase also encoded by the cassette. The EPSPS locus is also duplicated to a lesser degree in other weed species that have developed glyphosate resistance including *Kochia scoparia* [110,111]. In *K. scoparia* an initial unequal crossing-over is hypothesised to have led to duplication of EPSPS followed by selection for further unequal crossing-over events. Such a scenario could also explain our observation for the UGT locus as ONT long-reads indicate only the two UGT genes have undergone gene copy number amplification versus dozens as for *A. palmeri*. The two *P. ovis* UGT genes are located close to the start of Chromosome 7 in the *P. ovis* genome (sheep and cattle versions) at positions 159,563 to 161,298 kb (*P. ovis* cattle-derived genome), potentially in the subtelomeric region. Subtelomeric regions are enriched for repetitive elements, which result in genomic instability and increases the potential for unequal crossovers [112–114]. In the common bean, *Phaseolus vulgaris*, pathogen resistance (R) genes located in subtelomeric regions are highly-duplicated as a result of their location with associated resistance phenotypes [115]. Subtelomeric regions are variable in length from 10s to 100s of kilobases across organisms [112] and are currently not defined for *P. ovis*. The two *P. ovis* UGT genes may therefore occur within or nearby this region of instability which we hypothesise could, at least partially explain the elevated copy numbers of these genes through unequal crossing-over. This remains for confirmation in future studies alongside further investigation of inter-individual variability in UGT copy number and age of the expansion of UGT copy numbers in *P. ovis*.

#### 4.3.2 Transposable element mediated increased copy number of a CYP gene

The constitutively over-expressed CYP in our resistant RNASeq (RES_exposed_ and RES_unexposed_) replicates was associated with increased copy number in the six Resistant populations. The amplified region has clean break points and evidence for at least five potential insertion locations of this locus was provided by discordant read mappings (Supplementary Figure S1). It was not possible to confirm whether gene translocation events alone explain elevated coverage as a low number of long-reads used to assemble the *P. ovis* cattle genome indicated gene duplication *in-situ* may also have occurred at this locus. A relationship between transposable element (TE) activity and detoxification pathway gene overexpression has been observed in many arthropods [116,117]. The specific modes of TE action, especially as expression of duplicated genes is not simply doubled [118] remain to be elucidated but may involve loss or gain of promoter repressor and enhancement elements [117]. Several fixed synonymous positions in Resistant populations versus the Susceptible population and stop codon in adjacent PsoOvis1B003189 were identified. If a causal relationship between this gene and ML resistance is subsequently demonstrated in functional genomic experiments, these variants could form the basis of a simple PCR test for this form of ML resistance. In parasitic mites, no involvement of cytochrome P450s in ML resistance has been identified to date, but *S. scabiei* utilises this enzyme family for the detoxification of pyrethroids [119]. However, in other non-parasitic arthropods, CYPs have been linked to ML resistance [120]. For example, in *T. urticae,* CYPs have been linked to abamectin resistance [121]. Three different CYP genes were more highly expressed in different abamectin resistant strains [104]. A recombinant protein derived from one of these three genes was capable of detoxifying abamectin to a less toxic metabolite [91]. A nuclear receptor underlies trans-driven increases in detoxification genes including CYPs [19]. Knockout of the CYP gene CYP9A186 gene reversed avermectin resistance in the army worm, *Spodoptera exigua*. Recombinant CYP9A186 was able to metabolise abamectin and emamectin benzoate *in vitro* [122]. CYP also play a role in ivermectin resistance in the human body louse *Pediculus humanus humanus* and the tick *Rhipicephalus microplus*, although ABC-transporters have a bigger impact on susceptibility in *R. microplus* than CYPs [88,123,124].

The putative transposable element encoding gene PsoOvis1B002032 which had zero expression in SUS RNASeq replicates but very high expression in all six RES replicates, had low genome sequencing coverage in the sheep-derived genome assembly and is not present in the cattle-derived genome geneset. This may reflect the difficulty in assembling active TEs correctly, even with the excellent genomic resources available to us. Its striking expression dynamics remain an interesting observation, albeit restricted to the RNASeq component of our analyses.

### 4.4. Limitations in our experimental design

Metabolic resistance often occurs through constitutive over-expression of a limited number of genes in association with strong selection of involved loci. Underlying changes in copy number variation, as observed here, also frequently correlate with metabolic gene overexpression. The expression and pooled genome sequencing methods we applied here are well-suited to identifying such resistance arising from pesticide usage against arthropod targets. Due to this and reference to myriad other arthropod pests and disease vectors, it is likely that other genes and resistance mechanisms are involved in ML resistance in *P. ovis* cattle mites. For example, soft selective sweeps are less easily identified than hard sweeps seen at CYP loci in many organisms. The malaria transmitting mosquito, *Anopheles funestus* provides an obvious such example in temporal data pre- and post-introduction of pyrethroids [125]. Target-site changes undergoing soft sweeps which may not result in expression differences to confer resistance would be harder to detect for example.

We were also restricted to a single Susceptible population because of the dynamics of ML resistance in the sampling region which did not help in inferring subtle associations. This is a common issue for arthropod resistance to pesticide studies. For example, for some species of mosquito pyrethroid susceptible populations are no longer present in nature requiring comparison with laboratory-maintained populations and crossing experiments. As *P. ovis* occurs in multiple hosts this issue may be somewhat avoided by sampling rabbit or deer mite populations that have not experienced the selection pressure of ML use facing *P. ovis* on cattle or sheep. Such an experimental design would require care to avoid confounding signals of selection with any host specific adaptation. Despite these issues through gene expression and associated copy number variation we identified very strong candidate genes for ML-resistance in cattle-derived *P. ovis* mites.

### 4.5. Shared or distinct bases of ML-resistance in *P. ovis* on sheep?

*Psoroptes ovis* is found in several mammals but is likely a single species, although there is significant intraspecific morphological variation between cattle- and sheep-derived *P. ovis* [126]. Indeed, there is variation in the gene content of the two genome assemblies and fewer variants were predicted against the *P. ovis* cattle genome assembly than the *P. ovis* sheep assembly with our cattle-derived datasets. The two assemblies are not independent however as the *P. ovis* cattle genome was scaffolded using the sheep genome as a reference, hence caution is required when interpreting genetic differences from these genomes. It remains therefore for further investigation as to whether the genetic basis of ML resistance in sheep *P. ovis* is shared with cattle *P. ovis* or has arisen independently.

### 4.6. Conclusions

Our study is the first exploration of possible pharmacokinetic changes underlying ML resistance in *P. ovis* using extensive genomic and expression datasets. The over-expressed and multi-copy number UDP-glycosyltransferase and CYP genes represent strong candidates for ML resistance in *P. ovis* mites. Further research is required to functionally characterise these findings in the context of ML resistance. RNAi and functional expression and inhibition of these enzymes *in vivo*, are good approaches for demonstrating causation of detoxification pathway genes like UGTs and CYPs [14]. It is unlikely that detoxification genes are the only loci involved in conferring ML resistance as we observed peaks in F_st_ between Susceptible and Resistant populations that we have not linked to function. The inclusion of more ML-resistant *P. ovis* populations from other regions and host species is essential to determining if more than one resistance mechanism is segregating in *P. ovis*. Recent reports of ML-resistant *P. ovis* mite populations in sheep in the UK will aid in this process and represent important sample sets for future diagnostic test development [7,12]. This will aid the design of resistance tracking and management strategies and inform the likelihood of cross-resistance against second-line organophosphate treatments and novel acaricides in development.

## Acknowledgements

This research has received funding from the European Union’s Horizon 2020 research and innovation programme under grant agreement No 824110 – “EASI-Genomics”. Institutional support to CNAG was from the Spanish Government, Ministry of Science, Innovation and Universities and Generalitat de Catalunya through the Departament de Recerca i Universitats and Departament de Salut. The authors acknowledge Research Computing at the James Hutton Institute for providing computational resources and technical support for the “UK’s Crop Diversity Bioinformatics HPC” [127] (BBSRC grants BB/S019669/1 and BB/X019683/1), use of which has contributed to the results reported within this paper.

## Supplementary Methods

Detailed explanation of the steps, programs used and evaluation process for the sheep- and cattle-derived genome assemblies and associated genome annotation used in this manuscript.

## Supplementary Tables

**Supplementary Table S1.** GluCl-44 and GluCl-280 Primers with Illumina Adapters: Locus specific primer sequence bolded, Ns underlined and forward and Reverse barcoded sequencing primers with index sequence bolded. Illumina adaptor oligonucleotide sequences were obtained from the Illumina Adapter Sequences document dated March 2020 (Oligonucleotide sequences © 2020 Illumina, Inc.).

**Supplementary Table S2.** Forward and reverse primer sequences for the validation, by qPCR, of the selected *P. ovis* target genes and housekeeping genes.

**Supplementary Table S3.** Detailed description of the PCR products intended for Deep Amplicon Sequencing with the DNA concentration measured after PCR and the sample origin. The target gene of the PCR-product is indicated by the name in the first column with PSO44 (GluCl-44) and PSO280 (GluCl-280). The origin of the mite population is either from beef cattle farms visited in [13]or sheep from the Moredun Research Institute (MRI). NA = not available.

**Supplementary Table S4.** Detailed description of the RNA samples used for RNA seq. Three SUS replicates were extracted from the susceptible Farm 2 in van Mol et al, 2020 [13] and mites were collected before treatment. RES_unexposed_ replicates were extracted from resistant Farm 5 and mites were collected before treatment. Replicates RES_exposed_ 1 to 3 were from the same resistant farm, but mites were collected after treatment of the cattle with ivermectin. The RNA yields are presented together with the quality (RIN; RNA integrity number). Samples RES_exposed_ 1 and RES_exposed_ 2 contained equal numbers of mites (*) from the first and second collection. Sample RES_exposed_ 3 contained 100 mites from the first collection and 50 from the second (**).

**Supplementary Table S5.** Whole genome pooled template sequencing read alignment results for alignment against the cattle- and sheep-derived *P. ovis* genomes. Alignment statistics were generated using the “samtools stats” tool [54].

**Supplementary Table S6.** Coverage per gene (DXX) for each whole genome pooled template sequenced population aligned to the cattle genome. Coverage was divided by median coverage across the genome to identify outliers (DXX.median).

**Supplementary Table S7.** Coverage per gene (DXX) for each whole genome pooled template sequenced population aligned to the sheep genome. Coverage was divided by median coverage across the genome to identify outliers (DXX.median).

**Supplementary Table S8.** BLAST results for eight nanopore long reads with more than one copy of the CYP, PsoOvis1B011549, and neighbouring gene PsoOvis1B003189 in the cattle-derived mite genome assembly data. Results are in BLAST output format “6”.

**Supplementary Table S9.** BLAST results the two duplicated UGT genes against nanopore long-reads used in the cattle-derived mite genome assembly data. Results are in BLAST output format “6”.

**Supplementary Table S10.** BLAST results the two duplicated UGT genes against nanopore long-reads used in the sheep-derived mite genome assembly data. Results are in BLAST output format “6”.

**Supplementary Table S11.** Pairwise, average and absolute difference F_st_ values per gene for all populations. The final column in bold is the mean difference in F_st_ between average F_st_ for SUS vs RES comparisons and RES vs RES comparisons used to create Figure 10.

**Supplementary Table S12.** Frequencies per population, genomic, and gene-level locations for variants of interest in UGT and cytochrome P450 genes and nonsynonymous mutations in regions of high |F_st_| not otherwise linked to resistance phenotypes.

**Supplementary Table S13.** Raw data for input to rmcorr qPCR versus RNASeq correlation analysis used to create Figure 6.

## Supplementary Figures

**Supplementary Figure S1.** IGV plots of read alignment (BAM) files showing putative insertion or transposition sites across the *P. ovis* cattle-derived genome for a transposable element linked to amplification of the CYP gene PsoOvis1B011549. Five possible positions at four locations in the genome are shown by excessive read coverage with discordant read-mapping of pairs with the PsoOvis1B011549 locus.

## Supplementary Files

**Supplementary File S1.** Exposed ML-resistant (RES_exposed_) versus susceptible (SUS) mite DESeq2 results for all genes. Ranked by adjusted p-value.

**Supplementary File S2.** Unexposed ML-resistant (RES_unexposed_) versus susceptible (SUS) mite DESeq2 results for all genes. Ranked by adjusted p-value.

**Supplementary File S3.** Exposed ML-resistant (RES_exposed_) versus unexposed (RES_unexposed_) ML-resistant mite DESeq2 results for all genes. Ranked by adjusted p-value.

**Supplementary File S4.** Exposed ML-resistant (RES_exposed_) versus susceptible (SUS) mite Sleuth results for all isoforms of all genes. Ranked by adjusted p-value.

**Supplementary File S5.** Unexposed ML-resistant (RES_unexposed_) versus susceptible (SUS) mite Sleuth results for all isoforms of all genes. Ranked by adjusted p-value.

**Supplementary File S6.** Exposed ML-resistant (RES_exposed_) versus unexposed (RES_unexposed_) ML-resistant mite Sleuth results for all isoforms of all genes. Ranked by adjusted p-value.

## Notes

### Competing Interest Statement

The authors have declared no competing interest.

